# SorCS1-mediated sorting in dendrites maintains neurexin axonal surface polarization required for synaptic function

**DOI:** 10.1101/552695

**Authors:** Luís F. Ribeiro, Ben Verpoort, Julie Nys, Kristel M. Vennekens, Keimpe D. Wierda, Joris de Wit

## Abstract

The pre- and postsynaptic membranes comprising the synaptic junction differ in protein composition. The membrane trafficking mechanisms by which neurons control surface polarization of synaptic receptors remain poorly understood. The sorting receptor SorCS1 is a critical regulator of trafficking of neuronal receptors, including the presynaptic adhesion molecule neurexin (Nrxn), an essential synaptic organizer. Here, we show that SorCS1 maintains a balance between axonal and dendritic Nrxn surface levels in the same neuron. Newly synthesized Nrxn1α traffics to the dendritic surface, where it is endocytosed. Endosomal SorCS1 interacts with the Rab11 effector protein Rab11FIP5/Rip11 to facilitate the transition of internalized Nrxn1α from early to recycling endosomes and bias Nrxn1α surface polarization toward the axon. In the absence of SorCS1, Nrxn1α accumulates in early endosomes and mis-polarizes to the dendritic surface, impairing presynaptic differentiation and function. Thus, SorCS1-mediated sorting in dendritic endosomes controls Nrxn axonal surface polarization required for proper synapse development and function.

## Introduction

Neurons are highly compartmentalized cells that need to maintain distinct membrane identities. This is especially clear at the synapse, where pre- and postsynaptic membranes differ dramatically in their protein composition [1]. Membrane trafficking mechanisms that organize and maintain the polarized distribution of receptor proteins during the formation, maturation, and plasticity of synapses are of key importance for the proper function of neural circuits [2, 3].

Neurexins (Nrxns), expressed from three genes as α-, β-, and γ-Nrxns, are essential presynaptic adhesion molecules that engage in a network of interactions with multiple pre- and postsynaptic extracellular ligands [4] and act in a cell type-specific manner to regulate synapse number, function, and plasticity [5–9]. Mutations in *NRXNs* have been associated with multiple neuropsychiatric disorders, especially schizophrenia and autism [10–12], underscoring the importance of Nrxns for circuit function. Despite their essential role, the molecular mechanisms that control presynaptic polarization and abundance of Nrxns are largely unknown [13]. Previous studies have shown that surface trafficking of Nrxns requires the C-terminal PDZ-binding domain [14, 15] and that surface mobility depends on synaptic activity and interaction with extracellular ligands [16, 17].

The vertebrate SorCS1-3 (Sortilin-related CNS expressed 1-3) proteins are sorting receptors belonging to the VPS10P (vacuolar protein sorting 10) family receptors that have emerged as major regulators of intracellular protein trafficking [18, 19]. SorCS proteins are required for the surface expression of glutamate receptors, neurotrophin receptors, transporters, and adhesion molecules [20–24]. These sorting receptors are thus essential for synaptic transmission and plasticity, but the mechanisms by which SorCS proteins sort their cargo are poorly understood. Using quantitative proteomics analysis, we recently established a critical role for SorCS1 in Nrxn surface trafficking [20]. While our previous observations suggested that SorCS1 might regulate endosomal trafficking of Nrxn, the intracellular pathway via which Nrxn is trafficked to synapses, the regulation thereof by SorCS1, and the relevance of SorCS1-mediated sorting of Nrxn and other axonal cargo proteins for synaptic development and function remained elusive.

Here, we use epitope-tagging to elucidate the subcellular localization of endogenous SorCS1 and Nrxn1α. We show that SorCS1 acts in dendritic endosomes to control a balance between the axonal and dendritic surface distribution of Nrxn1α in the same neuron. Although predominantly considered to be a presynaptic molecule, we find that newly synthesized Nrxn1α first traffics to dendrites, where it is endocytosed, followed by transcytosis to the axon. SorCS1 controls Nrxn axonal surface polarization by facilitating the transition of endocytosed Nrxn from early to recycling endosomes and does so by interacting with Rab11FIP5/Rip11, a novel SorCS1 cytoplasmic binding partner we identify. In the absence of SorCS1, Nrxn1α accumulates in early endosomes and mis-polarizes to the dendritic surface, impairing presynaptic differentiation induced by the postsynaptic Nrxn ligand neuroligin. This defect can be rescued by a Nrxn1α mutant that bypasses SorCS1-mediated sorting and transcytosis to polarize to the axonal surface, but not by wildtype (WT) Nrxn1α. Finally, we show that SorCS1 is required for presynaptic function. Together, our observations indicate that SorCS1-mediated sorting in dendritic endosomes controls Nrxn axonal surface polarization required for proper synapse development and function.

## Results

### SorCS1 controls an axonal/dendritic balance in Nrxn1α surface polarization

We first determined the subcellular distribution of endogenous SorCS1, which has been difficult to assess due to a lack of suitable antibodies. We generated a SorCS1 knock-in (KI) mouse (*Sorcs1^HA^*), using CRISPR/Cas9 to insert a hemagglutinin (HA) epitope tag in the *Sorcs1* locus after the pro-peptide domain (Figure S1A). Correct insertion of the HA tag was confirmed by sequencing and western blot analysis (Figure S1B). HA immunodetection in cultured *Sorcs1^HA^* cortical neurons revealed prominent punctate endogenous SorCS1 immunoreactivity in soma and dendrites (Figure 1A), which was mimicked by exogenously expressed HA-SorCS1 in hippocampal neurons (Figure 1B, C). These results are in agreement with a previous immunohistochemical study showing a somatodendritic distribution of SorCS1 [25] and suggest that SorCS1-mediated sorting of cargo proteins, including Nrxn, occurs in this cellular compartment.

**Figure 1.**
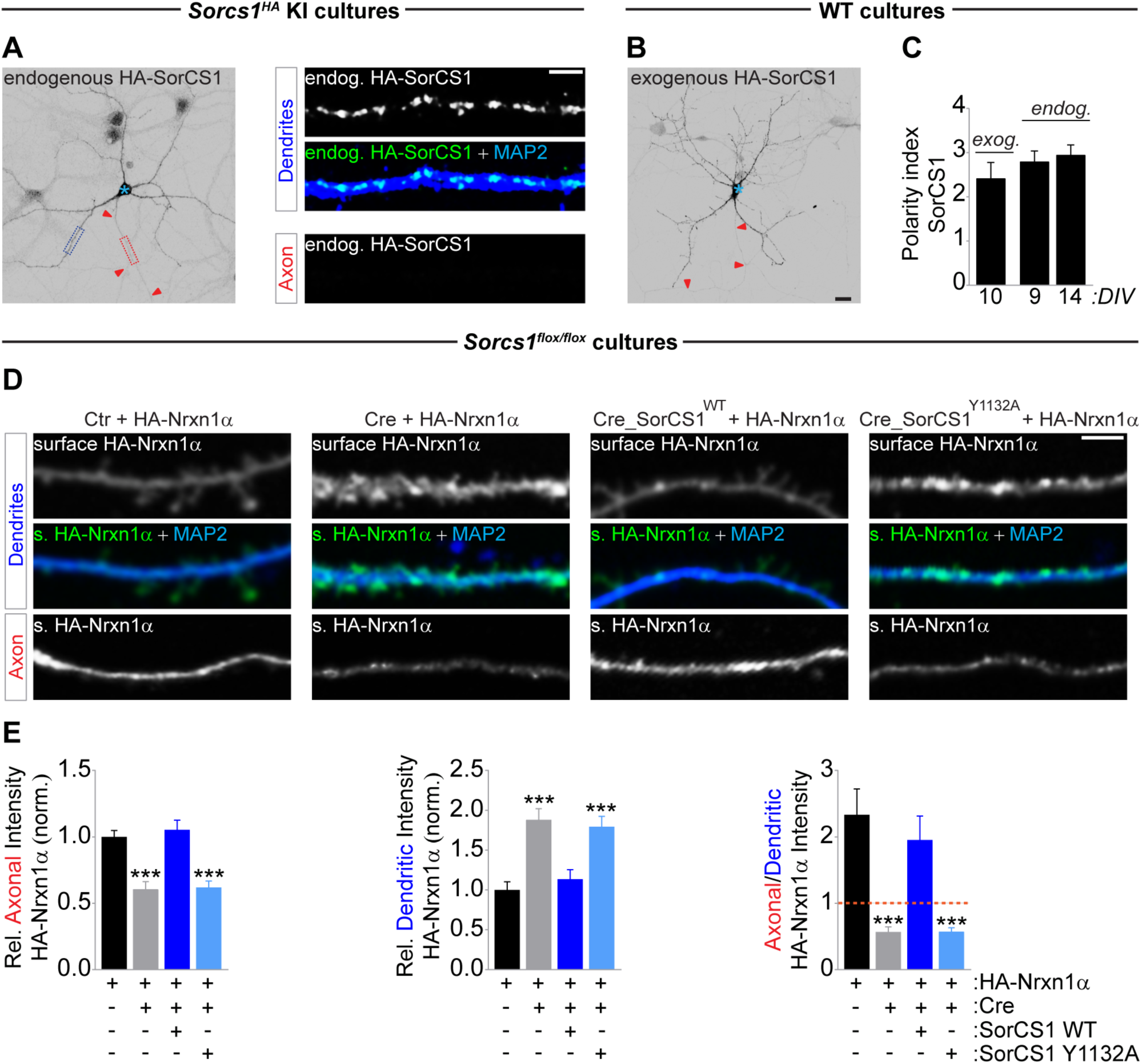
SorCS1-mediated sorting controls an axonal/dendritic surface balance of Nrxn1α. A, DIV9 *Sorcs1^HA^* mouse cortical neurons immunostained for endogenous (endog.) HA-SorCS1 (grayscale and green), MAP2 (blue) and Ankyrin-G (not shown). Red arrowheads indicate the axon and blue asterisk marks the cell body. High-zoom images show dendritic (dotted blue box) and axonal (dotted red box) distribution of HA-SorCS1. B, DIV10 WT mouse hippocampal neurons transfected with HA-SorCS1 and immunostained for HA (grayscale). MAP2 and Ankyrin-G not shown. C, Quantification of (A) and (B): dendritic versus axonal distribution (D:A – polarity index) of endogenous (DIV9, n = 26 neurons and DIV14, n = 26) and exogenous (exog.) (n = 14) HA-SorCS1 from 3 and 2 independent experiments, respectively. D, DIV8–DIV10 *Sorcs1^flox/flox^* mouse cortical neurons electroporated with EGFP (Ctr), Cre-EGFP, Cre-EGFP-T2A-SorCS1^WT^ or Cre-EGFP-T2A-SorCS1^Y1132A^ and transfected with HA-Nrxn1α, immunostained for surface (s.) HA-Nrxn1α (grayscale and green), MAP2 (blue); Ankyrin-G and GFP (not shown). E, Quantification of (D): surface HA-Nrxn1α fluorescence intensity in axon and dendrites relative to total surface levels and normalized to cells expressing EGFP, and ratio of axonal/dendritic surface HA intensity. Ctr (n = 28 neurons); Cre (n = 29); Cre_SorCS1^WT^ (n = 28); Cre_SorCS1^Y1132A^ (n = 28). ***p < 0.001 (Kruskal-Wallis test followed by Dunn’s multiple comparisons test, 3 independent experiments). Graphs show mean ± SEM. Scale bars, 20 µm (A, B); 5 µm (D).

To begin to elucidate how SorCS1 regulates membrane trafficking of its cargo Nrxn, we electroporated cortical *Sorcs1^flox/flox^*mouse neurons with Cre recombinase (Cre) to remove SorCS1 (*Sorcs1* KO). We transfected neurons with HA-Nrxn1α, one of the most abundant Nrxn isoforms in the brain [26], and live-labeled them with an HA antibody to detect surface Nrxn1α, followed by Ankyrin-G and MAP2 immunodetection to label the axonal and dendritic compartment, respectively. Loss of SorCS1 decreased Nrxn1α axonal surface levels and concomitantly increased dendritic surface levels in the same cell, changing Nrxn1α surface polarization from axonal to dendritic (Figure 1D, E). Re-expression of WT SorCS1 (SorCS1^WT^) in Cre-positive *Sorcs1^flox/flox^*neurons restored Nrxn1α axonal surface polarization (Fig. 1D, E), indicating that the defect occurred cell-autonomously. In contrast, rescue with an endocytosis-defective mutant [27] of SorCS1 (SorCS1^Y1132A^), which remains on the plasma membrane (Figure S1C, D), did not rescue Nrxn1α surface polarization (Figure 1D, E). To determine whether SorCS1 controls the subcellular distribution of endogenous Nrxn, we immunostained *Sorcs1^flox/flox^*neurons electroporated with Cre or GFP with a pan-Nrxn antibody directed against the conserved cytoplasmic tail [28] (Figure S1E, F), which specifically labels endogenous Nrxn (Figure S1G, H). Similar to the observations with transfected HA-Nrxn1α in *Sorcs1* KO neurons, loss of SorCS1 reduced axonal intensity of endogenous Nrxn and decreased the axonal/dendritic ratio compared to control neurons (Figure S1E, F). Conversely, overexpression of SorCS1^WT^ in hippocampal neurons, which express very low levels of *Sorcs1* [29], increased Nrxn1α axonal surface polarization and concurrently decreased dendritic surface levels, whereas overexpression of SorCS1^Y1132A^ did not alter Nrxn1α surface polarization (Figure S1I, J). Together, these results indicate that somatodendritic SorCS1 controls an axonal/dendritic surface balance of Nrxn1α. Our findings suggest that SorCS1 is either required for endocytosis of Nrxn1α or acts in endosomes to bias Nrxn1α surface polarization toward the axon.

### Indirect axonal surface trafficking of Nrxn1α in mature neurons

Nrxns are considered to be predominantly presynaptic proteins, although several independent studies have reported the presence of a dendritic pool of Nrxns [20,30–33]. To unequivocally determine the subcellular localization of endogenous Nrxn, we generated a Nrxn1α KI mouse (*Nrxn1α^HA^*), inserting an HA epitope tag in the *Nrxn1α* locus after the signal peptide (Figure S2A). Correct insertion of the HA tag was confirmed by restriction digest, sequencing, and western blot analysis (Figure S2B). We cultured *Nrxn1α^HA^* cortical neurons together with WT cortical neurons to reliably detect endogenous HA-Nrxn1α in the somatodendritic and axonal compartment (Figure S2C, D). At 3 days in vitro (DIV), HA-Nrxn1α surface distribution was polarized toward the axon (Figure 2A, B). As neurons matured, surface HA-Nrxn1α axonal surface levels decreased, while dendritic surface levels increased. Surface HA-Nrxn1α remained polarized towards the axon at all developmental time points analyzed (Figure 2A, B). A similarly polarized distribution of Nrxn was observed in WT neurons labeled with the pan-Nrxn antibody (Figure S2E, F). Together, these results indicate that endogenous Nrxn1α is an axonally polarized surface protein that accumulates in dendrites as neurons mature.

**Figure 2.**
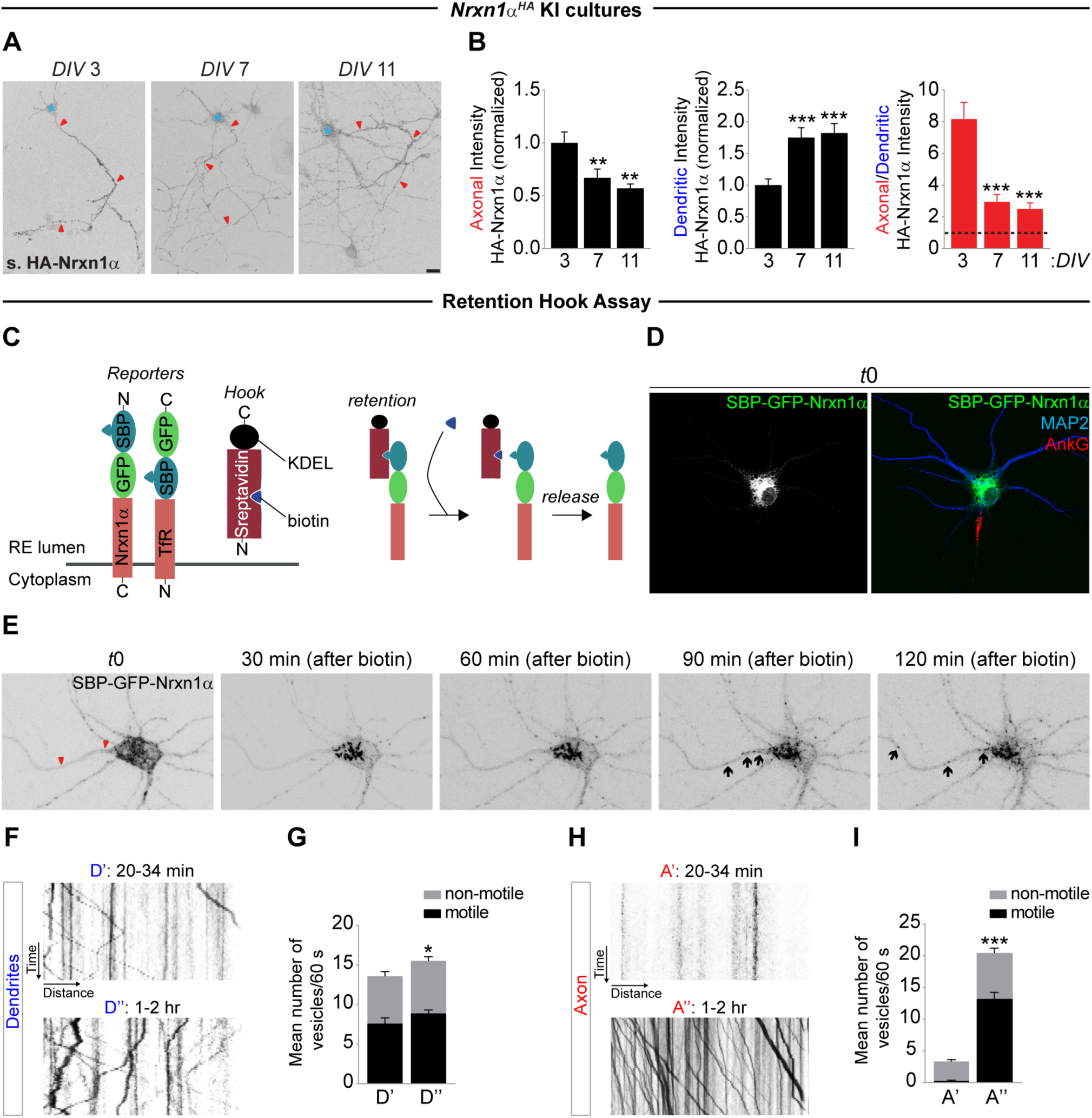
Indirect axonal trafficking of Nrxn1α in mature neurons. A, Non-permeabilized DIV3, DIV7 and DIV11 *Nrxn1α^HA^* cortical neurons live-labeled with an HA antibody to visualize endogenous surface (s.) HA-Nrxn1α in axon and dendrites. Red arrowheads indicate the axon and blue asterisk marks the cell body. B, Quantification of (A): surface HA-Nrxn1α fluorescence intensity in axon and dendrites normalized to DIV3 neurons, and ratio of axonal/dendritic surface HA-Nrxn1α intensity. DIV3 (n = 30 neurons); DIV7 (n = 29); DIV11 (n = 24). **p < 0.01; ***p < 0.001 (Kruskal-Wallis test followed by Dunn’s multiple comparisons test, 3 independent cultures). C, Schematic representation of streptavidin-KDEL (ER hook) and reporters (Nrxn1α and TfR) used in RUSH experiments. Addition of biotin dissociates the reporters from the ER hook, inducing synchronous release from the ER and transport through the secretory pathway. D, DIV9 WT mouse cortical neuron co-expressing SBP-GFP-Nrxn1α and streptavidin-KDEL immunostained for MAP2 (blue), Ankyrin-G (red) and GFP-Nrxn1α (grayscale and green) at t=0 before adding biotin. E, Live-cell imaging in DIV8–DIV10 WT rat cortical neurons co-expressing SBP-GFP-Nrxn1α and ER hook. After 24–31 hr of expression neurons were imaged every 5 min for 2.5 hr. Biotin was added 10 min after the beginning of the imaging session. See also Supplementary Movie 1. F, Kymographs illustrating GFP-Nrxn1α vesicle dynamics over a 60s period in dendrites [D] from neurons treated with biotin for either 25 min or 2 hr. See also Supplementary Movie 2 and 3. G, Mean number of motile and non-motile GFP-Nrxn1α vesicles in dendrites from neurons treated with biotin for either 20–34 min (n = 17 neurons) or 1–2 hr (n = 16) in 2 and 3 independent experiments, respectively. H, Kymographs illustrating GFP-Nrxn1α vesicle dynamics over a 60s period in axons [A] from neurons treated with biotin for either 25 min or 2 hr. See also Supplementary Movie 2 and 3. I, Mean number of motile GFP-Nrxn1α vesicles in axons. Graphs show mean ± SEM. *p < 0.05; **p < 0.01; ***p < 0.001 (Mann-Whitney test). Scale bars, 20 µm (A, E); 10 µm (D).

To determine how Nrxn1α is trafficked in cortical neurons, we employed the retention using selective hooks (RUSH) system [34] to retain Nrxn1α in the endoplasmic reticulum (ER) and induced synchronous release and transport through the secretory pathway by applying biotin (Figure 2C). We performed live-cell imaging in 8–10 days in vitro (DIV) rat cortical neurons co-expressing SBP-GFP-Nrxn1α (reporter) and streptavidin-KDEL (ER hook), which were live-labeled with an antibody against the axon initial segment protein Neurofascin to visualize the axonal compartment. At t=0, we observed diffuse SBP-GFP-Nrxn1α fluorescence in soma and dendrites, in a pattern resembling the ER (Figure 2D). Following application of biotin, Nrxn1α fluorescence coalesced into punctate structures that started trafficking abundantly in the somatodendritic compartment, followed by Nrxn1α trafficking in the axon after a marked delay (Figure 2E; Movie S1). Quantification of Nrxn1α fluorescence intensity showed a decrease in Nrxn1α intensity in the soma and an increase in axonal intensity approximately 90 min after biotin application (Figure S3A). Kymograph analysis showed a small increase in the mean number of motile vesicles present in dendrites at late (1–2 hr) compared to early (20–34 min) time-points (Figure 2E, F; Movies S2, S3). Shortly after ER release, the majority of vesicles in dendrites moved in the anterograde direction, shifting to a retrograde movement at the later time-point (Figure S3B). Strikingly, motile Nrxn1α vesicles were absent in axons shortly after ER release but increased dramatically at late time-points (Figure 2G, H, Movies S2, 3). The vast majority of these vesicles moved anterogradely (Figure S3C).

We performed several controls to verify that the delayed axonal trafficking of Nrxn1α is not an artefact of prolonged retention in the ER and synchronized release, which might overwhelm the trafficking machinery. Trafficking of the somatodendritic cargo protein TfR [35] was detected only in the somatodendritic compartment (Figure S3D, E; Movie S4). In the absence of biotin, dendritic SBP-GFP-Nrxn1α-positive structures were not motile (Movie S5), indicating that the delayed trafficking of Nrxn1α to axons is not simply caused by prolonged ER retention. Moreover, immature DIV3 neurons displayed Nrxn1α trafficking to the axonal compartment shortly after biotin application (Figure S3F-H; Movie S6), further indicating that the delayed axonal trafficking of Nrxn1α in mature neurons does not result from prolonged retention in the ER but correlates with the maturation state of the neuron. Taken together, these results show that following transport through the secretory pathway, Nrxn1α is first trafficked to the somatodendritic compartment and appears in axons with a marked delay.

### Nrxn1α is transcytosed from the dendritic surface to axons

Previous work has shown that axonal polarization of membrane proteins via indirect trafficking can be achieved by non-polarized delivery and selective retention in the axon, or by indirect polarized delivery via insertion in dendrites followed by transcytosis to the axon [36]. To determine via which indirect pathway Nrxn1α is trafficked to the axonal surface, we followed the fate of Nrxn1α throughout the endosomal pathway. Internalized Nrxn1α was largely restricted to the dendritic compartment (Figure S4A, B). Blocking of dynamin-dependent endocytosis, either by expressing a dominant-negative mutant of Dynamin1 (K44A) or with Dynasore (Figure S4C-F), caused a decrease in Nrxn1α axonal surface levels and a concomitant increase in dendritic surface levels. Expression of GTP binding-deficient mutants of Rab5 (S34N) and Rab11 (S25N), to interfere with transport to early and recycling endosomes, respectively, mimicked the effect of blocking endocytosis (Figure S4G–J). However, expression of Rab7 (T22N) to interfere with transport to late endosomes did not affect the axonal surface polarization of Nrxn1α (Figure S4K, L). Thus, endocytosis, early and recycling endosomal transport, but not late endosomal transport, are required for accumulation of Nrxn1α on the axonal plasma membrane, indicating transcytotic trafficking of Nrxn1α.

### A SorCS1-Rab11FIP5 interaction controls Nrxn1α transition from early to recycling endosomes

The transcytotic trafficking route of Nrxn1α and somatodendritic distribution of SorCS1 suggest that sorting of Nrxn1α occurs in dendrites. To determine how SorCS1 controls transcytotic trafficking of Nrxn1α, we followed the fate of internalized HA-Nrxn1α in *Sorcs1* KO neurons. Endocytosis of HA-Nrxn1α in dendrites was not affected in *Sorcs1* KO neurons compared to control cells (Figure S5A, B). We next analyzed colocalization of internalized Nrxn1α with EEs (EAA1), fast REs (Rab4), REs (Rab11) in dendrites, and REs (Rab11) in the axon, respectively of *Sorcs1* KO neurons. Quantification revealed an increase in the colocalization of internalized Nrxn1α with EAA1-positive early endosomes (EEs) and Rab4-positive fast recycling endosomes (REs) in dendrites of *Sorcs1* KO neurons (Figure 3A-D). Colocalization of internalized Nrxn1α with Rab11-positive REs in *Sorcs1* KO neurons was decreased in dendrites (Figure 3E, F) and even more strongly reduced in the axon (Figure 3G, H). Thus, endocytosed Nrxn1α accumulates in EEs and is mis-sorted to Rab4-fast REs in the absence of SorCS1-mediated sorting. These observations are consistent with our finding that dendritic surface levels of Nrxn1α are increased in *Sorcs1* KO neurons (Figure 1D, E), which is likely due to increased recycling of Nrxn1α via Rab4-fast REs back to the dendritic plasma membrane. Similarly, the reduced axonal surface levels of Nrxn1α in *Sorcs1* KO neurons (Figure 1D, E) likely result from decreased sorting of Nrxn1α to Rab11-REs. Together, these results demonstrate that SorCS1 controls axonal surface polarization of Nrxn1α by facilitating the transition from EEs to Rab11-REs.

**Figure 3.**
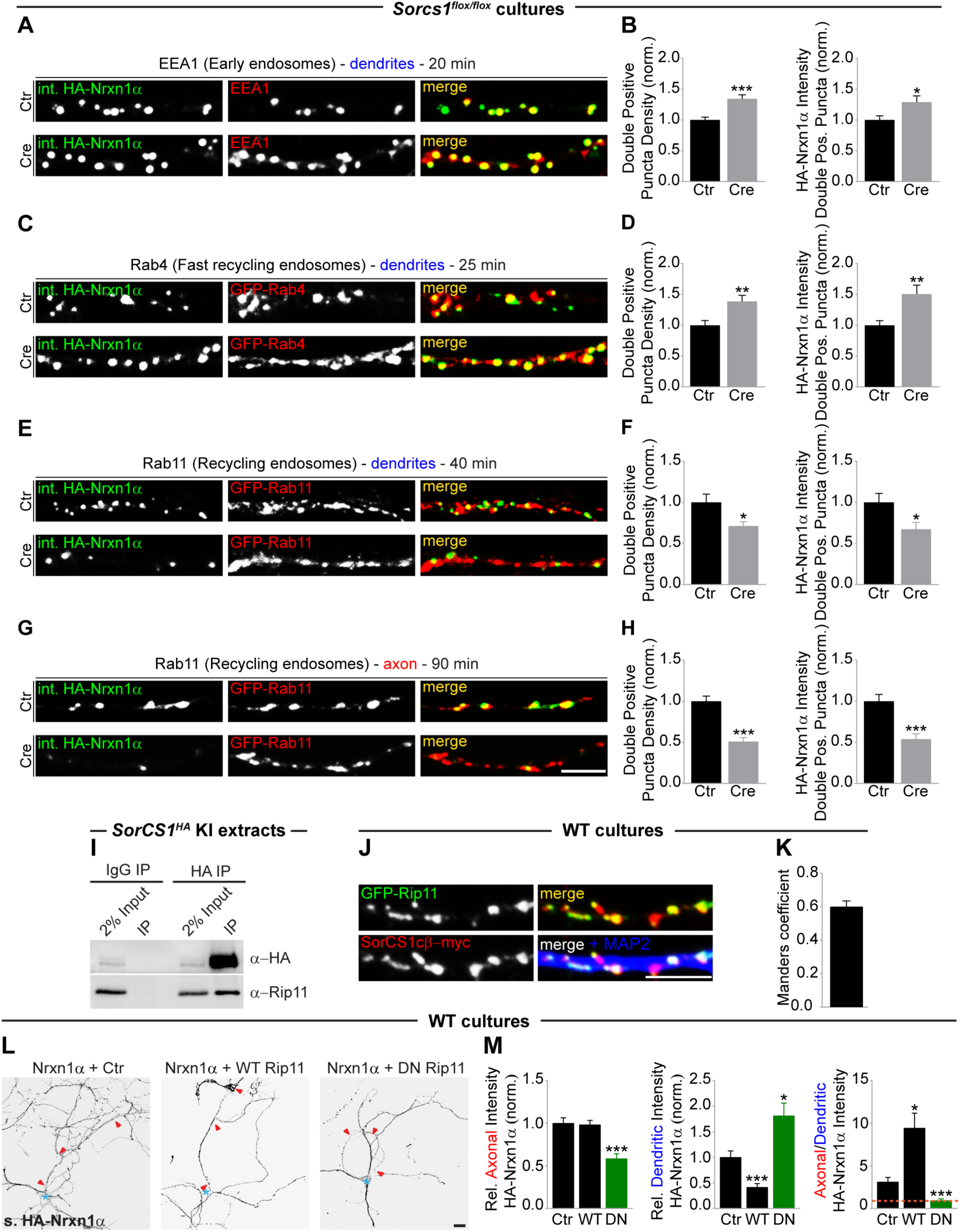
SorCS1 interacts with Rab11FIP5/Rip11 to regulate Nrxn1α transition from early to recycling endosomes. A, DIV8–DIV10 *Sorcs1^flox/flox^* cortical neurons electroporated with mCherry (Ctr) or Cre-T2A-mCherry and transfected with HA-Nrxn1α (pulse-chased for 20 min), labeled for internalized (int.) HA-Nrxn1α (grayscale), EEA1 (red) and Ankyrin-G (not shown). B, Quantification of (A): number of internalized Nrxn1α- and EEA1-double positive puncta normalized to cells expressing mCherry, and intensity of internalized Nrxn1α fluorescence in the double-positive puncta normalized to cells expressing mCherry. Ctr (n = 26 neurons); Cre (n = 25). *p < 0.05; ***p < 0.001 (Mann-Whitney test, 3 independent experiments). C, DIV8–DIV10 *Sorcs1^flox/flox^* cortical neurons electroporated with mCherry (Ctr) or Cre-T2A-mCherry and co-transfected with HA-Nrxn1α and GFP-Rab4 (pulse-chased for 25 min), labeled for internalized HA-Nrxn1α (grayscale), GFP-Rab4 (red) and Ankyrin-G (not shown). D, Quantification of (C); Ctr (n = 28 neurons); Cre (n = 26). **p < 0.01 (Mann-Whitney test, 3 independent experiments). E, DIV8–DIV10 *Sorcs1^flox/flox^* cortical neurons electroporated with mCherry (Ctr) or Cre-T2A-mCherry and co-transfected with HA-Nrxn1α and GFP-Rab11 (pulse-chased for 40 min), labeled for internalized HA-Nrxn1α (grayscale), GFP-Rab11 (red) and Ankyrin-G (not shown). F, Quantification of (E); Ctr (n = 29 neurons); Cre (n = 23). *p < 0.05 (Mann-Whitney test, 3 independent experiments). G, DIV8–DIV10 *Sorcs1^flox/flox^* cortical neurons electroporated with mCherry (Ctr) or Cre-T2A-mCherry and co-transfected with HA-Nrxn1α and GFP-Rab11 (pulse-chased for 90 min), labeled for internalized HA-Nrxn1α (grayscale), GFP-Rab11 (red) and Ankyrin-G (not shown). H, Quantification of (G); Ctr (n = 28 neurons); Cre (n = 26). ***p < 0.001 (Mann-Whitney test, 3 independent experiments). I, Western blot for the recovery of Rab11FIP5/Rip11 in immunoprecipitated HA-SorCS1 complexes from P21–P28 *Sorcs1^HA^* cortical prey extracts. J, DIV9–DIV10 WT cortical neurons co-expressing EGFP-Rip11 and SorCS1cβ-myc immunostained for GFP-Rip11 (grayscale and green), SorCS1-myc (grayscale and red) and MAP2 (blue). K, Quantification of the colocalization of Rip11 with SorCS1 expressed as Manders coefficient (n = 20 neurons) in 2 independent experiments. L, DIV9 WT cortical neurons co-expressing HA-Nrxn1α and EGFP (Ctr), WT EGFP-Rip11 or dominant-negative (DN) EGFP-Rip11, and immunostained for surface (s.) HA-Nrxn1α (grayscale); MAP2, Ankyrin-G and GFP (not shown). Red arrowheads indicate the axon and blue asterisk marks the cell body. M, Quantification of (L): surface HA-Nrxn1α fluorescence intensity in axon and dendrites relative to total surface levels and normalized to cells expressing EGFP and ratio of axonal/dendritic surface HA intensity (n = 30 for each group). *p < 0.05; ***p < 0.001 (Kruskal-Wallis test followed by Dunn’s multiple comparisons test, 3 independent experiments). Graphs show mean ± SEM. Scale bars, 5 µm (A, C, E, G, J); 20 µm (L).

We reasoned that SorCS1 might interact with additional proteins to facilitate Nrxn1α sorting from early to recycling endosomes. Rab11 family-interacting protein 5 (Rab11FIP5/Rip11 or Rip11), is prominently present in the raw mass spectrometric (MS) data set we obtained after affinity purification (AP) of SorCS1 complexes from rat brain extracts with two independent antibodies [20] (AP-MS data available online). Rip11, which belongs to the family of Rab11-interacting proteins [37], localizes to REs and regulates transcytosis of proteins from the basolateral to the apical plasma membrane in polarized epithelial cells [38, 39]. Western blot analysis of immunoprecipitated HA-SorCS1 from postnatal *Sorcs1^HA^* KI cortical extracts showed a robust Rip11 band coprecipitating with SorCS1, but not with control mouse IgG (Figure 3I). Rip11 displayed a punctate distribution in dendrites and colocalized with SorCS1 (Figure 3J, K). Expression of a dominant-negative (DN) form of Rip11 that inhibits the transport from early to recycling endosomes [40] reduced axonal surface levels of HA-Nrxn1α and increased dendritic surface levels, shifting Nrxn1α surface polarization from axonal to dendritic (Figure 3L, M), similar to loss of SorCS1. Expression of WT Rip11 on the other hand increased axonal surface polarization of HA-Nrxn1α (Figure 3L, M). Together, these results demonstrate that a SorCS1-Rip11 interaction facilitates Nrxn1α sorting from early to recycling endosomes, preventing mis-sorting to fast recycling and late endosomes and biasing trafficking to the axonal surface.

### Selective mis-sorting of transcytotic cargo in the absence of SorCS1

We next asked whether the surface polarization of other axonal membrane proteins was affected in the absence of SorCS1. The cell adhesion molecules L1/NgCAM and Caspr2 are both targeted to the axonal surface, but via distinct trafficking routes. L1/NgCAM is transcytosed from dendrites to axon [41, 42], whereas Caspr2 undergoes non-polarized delivery followed by selective endocytosis from the somatodendritic surface resulting in axonal polarization [43]. L1/NgCAM levels are downregulated in *Sorcs1* KO synaptosomes [20]. *Sorcs1^flox/flox^* cortical neurons electroporated with Cre or GFP were transfected with myc-L1 or HA-Caspr2 and immunostained for surface myc or HA. Consistent with a role for SorCS1 in regulating transcytosed axonal cargo, surface polarization of L1/NgCAM, but not of Caspr2, was perturbed in *Sorcs1* KO neurons (Figure S5C–E). The polarity index of two somatodendritic proteins (GluA2-GFP and MAP2), was unchanged in *Sorcs1* KO neurons (data not shown), indicating that loss of SorCS1 does not generally perturb the polarized distribution of neuronal proteins. Thus, loss of SorCS1 selectively impairs transcytotic trafficking of axonal membrane proteins, while keeping other neuronal polarity mechanisms intact.

### Loss of SorCS1 impairs Nrxn-mediated presynaptic differentiation

We next sought to determine motifs in the Nrxn1α protein sequence that contain signals necessary for its membrane trafficking, with the goal of identifying a Nrxn1α mutant that can bypass SorCS1-mediated sorting and transcytotic trafficking. We systematically tested the effect of a series of Nrxn1α cytoplasmic deletions on surface polarization (data not shown) and found that removing the 4.1-binding motif in the cytoplasmic domain (Nrxn1α Δ4.1) was the only deletion that dramatically increased Nrxn1α axonal surface polarization (Figure 4A, B). Remarkably, axonal surface polarization of Nrxn1α Δ4.1 was unaffected by loss of SorCS1 (Figure 4C, D). Moreover, Dynasore treatment to block endocytosis did not impair axonal surface polarization of Nrxn1α Δ4.1 (Figure 4E, F), indicating that the Nrxn1α Δ4.1 mutant bypasses dendritic endocytosis and sorting by SorCS1 to polarize to the axonal surface.

**Figure 4.**
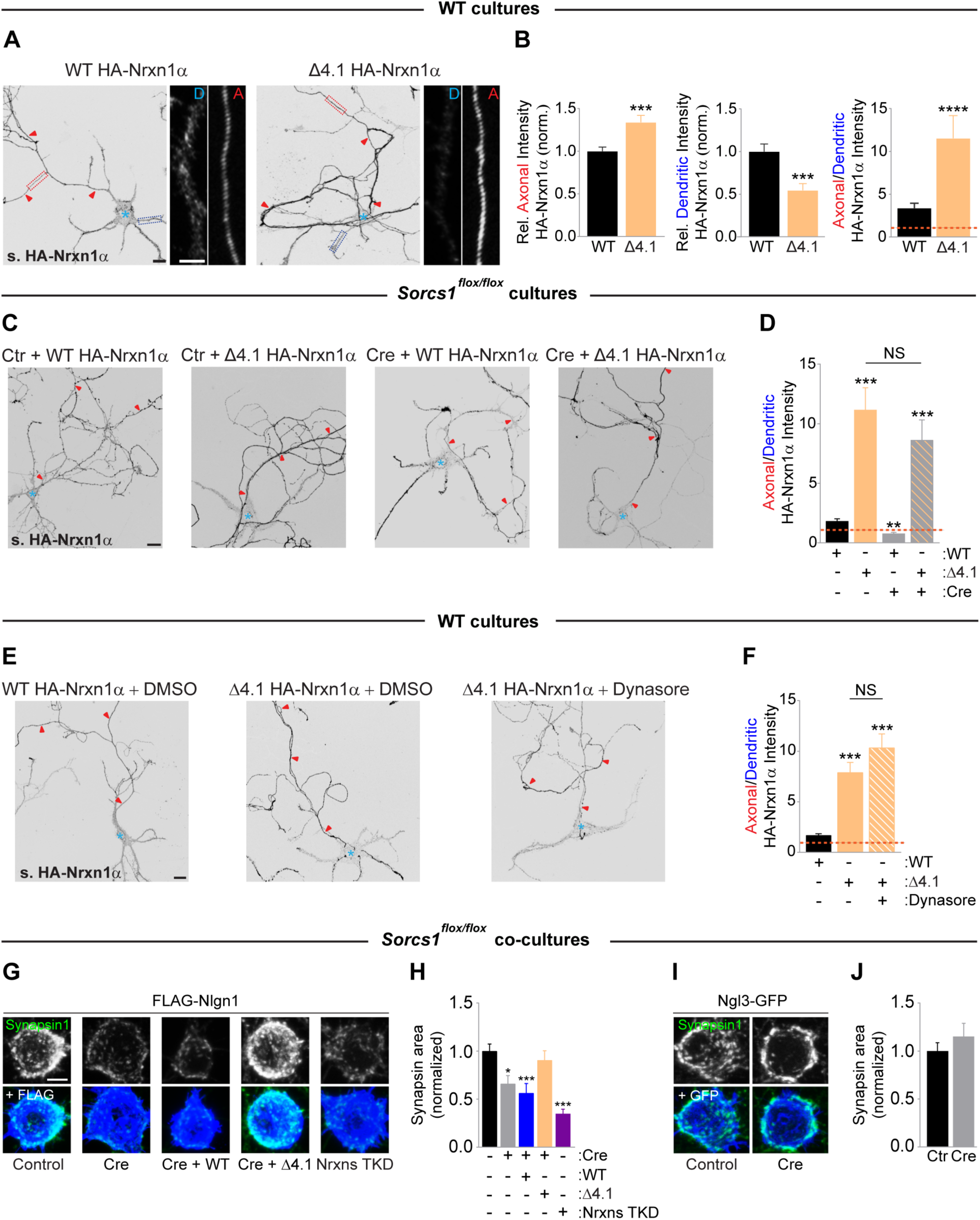
SorCS1-mediated axonal surface polarization of Nrxn is required for presynaptic differentiation. A, DIV8–DIV10 WT mouse cortical neurons transfected with WT HA-Nrxn1α and a cytoplasmic deletion mutant lacking the 4.1-binding motif (HA-Nrxn1α Δ4.1) and immunostained for surface (s.) HA-Nrxn1α (grayscale); MAP2 and Ankyrin-G (not shown). Red arrowheads indicate the axon and blue asterisk marks the cell body. High-zoom images show dendritic [D, dotted blue box] and axonal [A, dotted red box] HA-Nrxn1α. B, Quantification of (A): surface HA-Nrxn1α fluorescence intensity in axon and dendrites relative to total surface levels and normalized to cells expressing WT-Nrxn1α and ratio of axonal/dendritic surface HA intensity. WT (n = 40 neurons); Δ4.1 (n = 30). ***p < 0.001 ****p < 0.0001 (Mann-Whitney test, 3 independent experiments). C, DIV8–DIV10 mouse *Sorcs1^flox/flox^* cortical neurons electroporated with EGFP (Ctr) or Cre-EGFP (Cre) and transfected with WT HA-Nrxn1α or HA-Nrxn1α Δ4.1. Neurons were immunostained for surface HA-Nrxn1α (grayscale); MAP2, Ankyrin-G and GFP (not shown). D, Quantification of (C): ratio of axonal/dendritic surface HA-Nrxn1α fluorescence intensity. Ctr_WT (n = 46 neurons); Ctr_Δ4.1 (n = 25); Cre_WT (n = 30); Cre_Δ4.1 (n = 27). NS, non-significant; **p < 0.01; ***p < 0.001 (Kruskal-Wallis test followed by Dunn’s multiple comparisons test, at least 3 independent experiments). E, DIV8–DIV10 WT mouse cortical neurons transfected with WT HA-Nrxn1α or HA-Nrxn1α Δ4.1 and treated with DMSO (vehicle) or Dynasore. Neurons were immunostained 18 hr after treatment for surface HA-Nrxn1α (grayscale); MAP2 and Ankyrin-G (not shown). F, Quantification of (E): ratio of axonal/dendritic surface HA fluorescence intensity. WT_DMSO (n = 40 neurons); Δ4.1_DMSO (n = 40); Δ4.1_Dynasore (n = 28). NS, non-significant; ***p < 0.001 (Kruskal-Wallis test followed by Dunn’s multiple comparisons test, at least 3 independent experiments). G, HEK293T-cells expressing FLAG-Nlgn1 co-cultured with DIV10 *Sorcs1^flox/flox^*cortical neurons infected with LV expressing mCherry (Control), Cre-T2A-mCherry, Cre-T2A-mCherry-T2A-WT Nrxn1α, Cre-T2A-mCherry-T2A-Nrxn1α Δ4.1 or Nrxn TKD; immunostained for FLAG (blue), Synapsin1 (green) and mCherry (not shown). H, Quantification of (G): the area of Synapsin1 clustering on the surface of Nlgn1-expressing HEK cells and normalized to cells expressing mCherry. Control (n = 29 neurons); Cre (n = 30); Cre-WT (n = 30); Cre-Δ4.1 (n = 30); TKD Nrxns (n = 30). *p < 0.05; ***p < 0.001 (Kruskal-Wallis test followed by Dunn’s multiple comparisons test, 3 independent experiments). I, HEK293T-cells expressing NGL-3-GFP co-cultured with DIV10 *Sorcs1^flox/flox^*cortical neurons infected with LV expressing mCherry (Control) or Cre-T2A-mCherry. J, Quantification of (I). Control (n = 30 neurons); Cre (n = 30). Graphs show mean ± SEM. Scale bars, 20 µm (A, C, E); 5 µm (A [high-zoom], G, I).

Nrxn mis-sorting to the dendritic surface and its concomitant loss from the axonal surface in *Sorcs1* KO neurons would be expected to impair synapse formation on heterologous cells expressing a postsynaptic ligand for Nrxn [44]. To test this, we infected *Sorcs1^flox/flox^* cortical neurons with lentivirus (LV) to express Cre-T2A-mCherry or mCherry as control and co-cultured these with HEK293T cells expressing the Nrxn ligand FLAG-Neuroligin 1 (Nlgn1). Nlgn1-induced clustering of the presynaptic marker Synapsin 1 was reduced in *Sorcs1* KO axons compared to control axons (Figure 4G, H). This defect was specific, as presynaptic differentiation induced by Netrin-G Ligand 3 (NGL-3), which requires presynaptic leukocyte common antigen-related protein (LAR) [45], was not affected in *Sorcs1* KO neurons (Figure 4I, J). Infection with a lentiviral vector expressing shRNAs against all Nrxns (Nrxn TKD) [46] mimicked the defect in *Sorcs1* KO neurons (Figure 4G, H), suggesting that Nlgn1-induced presynaptic differentiation is impaired in *Sorcs1* KO neurons due to decreased axonal surface levels of Nrxn. To test this, we overexpressed HA-Nrxn1α in *Sorcs1* KO neurons using LV. As expected, WT Nrxn1α did not rescue the impaired Nlgn1-mediated synaptogenic activity (Figure 4G, H), due to Nrxn1α mis-polarization to the dendritic surface in *Sorcs1* KO neurons. In contrast, expression of the Nrxn1α Δ4.1 mutant, which bypasses SorCS1-mediated sorting and the transcytotic route to polarize to the axonal surface, rescued the synaptogenic defect caused by SorCS1 loss (Figure 4G, H). Together, these results show that SorCS1-mediated sorting of Nrxns, by promoting Nrxn accumulation on the axonal surface, is required for normal synaptogenesis onto Nlgn1-expressing cells.

### SorCS1-mediated sorting is required for presynaptic function

Mis-sorting of Nrxns, which regulate neurotransmitter release [5–7], would also be expected to impair presynaptic function. To assess the consequences of loss of SorCS1 on synaptic function, we recorded spontaneous miniature excitatory postsynaptic currents (mEPSCs) from *Sorcs1^flox/flox^*autaptic cortical cultures electroporated with Cre or GFP. SorCS1 loss strongly decreased mEPSC frequency (Figure 5A, B). Decay kinetics and amplitude of mEPSCs were not altered by loss of SorCS1 (Figure 5B), suggesting that the amount of neurotransmitter release per vesicle and postsynaptic receptor properties and/or receoptor density were not affected. The amplitude and total charge transfer of single evoked EPSCs (eEPSCs) were reduced in *Sorcs1* KO neurons (Figure 5C, D). The mEPSC frequency and eEPSC amplitude defects can be attributed to a decrease in the readily releasable pool (RRP) size, in the vesicular release probability (Pves), or in active synapse number. Indeed, we previously found an increase in the number of silent excitatory synapses after loss of SorCS1 [20]. To assess RRP size, we applied a hyperosmotic sucrose stimulus. The amplitude and total charge transfer of synaptic responses (Figure 5E, F) and RRP size (Figure 5G) induced by single application of 0.5 M sucrose were strongly decreased in *Sorcs1* KO neurons, in line with a reduction in active synapse number. Because eEPSC amplitude and RRP size were proportionally reduced in *Sorcs1* KO cells, the Pves (evoked EPSC charge/initial sucrose charge) was unchanged (Figure 5H). To further examine potential presynaptic release defects in *Sorcs1* KO neurons, we performed train stimulations at different frequencies to probe for short-term plasticity defects that cannot be detected by induction of single EPSC and single sucrose application. Repeated stimulation at 10 Hz produced a more pronounced rundown of normalized evoked responses (synaptic depression) in *Sorcs1* KO neurons compared to control cells (Figure 5I, J), suggesting a presynaptic defect during sustained periods of activity. Other stimulation frequencies and paired-pulse stimulations also showed increased synaptic depression in *SorCS1* KO neurons (data not shown). Calculating the RRP size by back-extrapolation of cumulative synchronous charge during 10 Hz-train stimulation confirmed a reduction in RRP size in *Sorcs1* KO neurons (Figure 5K), indicating a presynaptic defect independent of the silent synapses phenotype that we described previously [20]. Together, these observations indicate that loss of SorCS1 impairs neurotransmitter release, reminiscent of Nrxn loss-of-function [5–7], indicating that SorCS1-mediated sorting of Nrxns in dendrites is required for normal presynaptic function.

**Figure 5.**
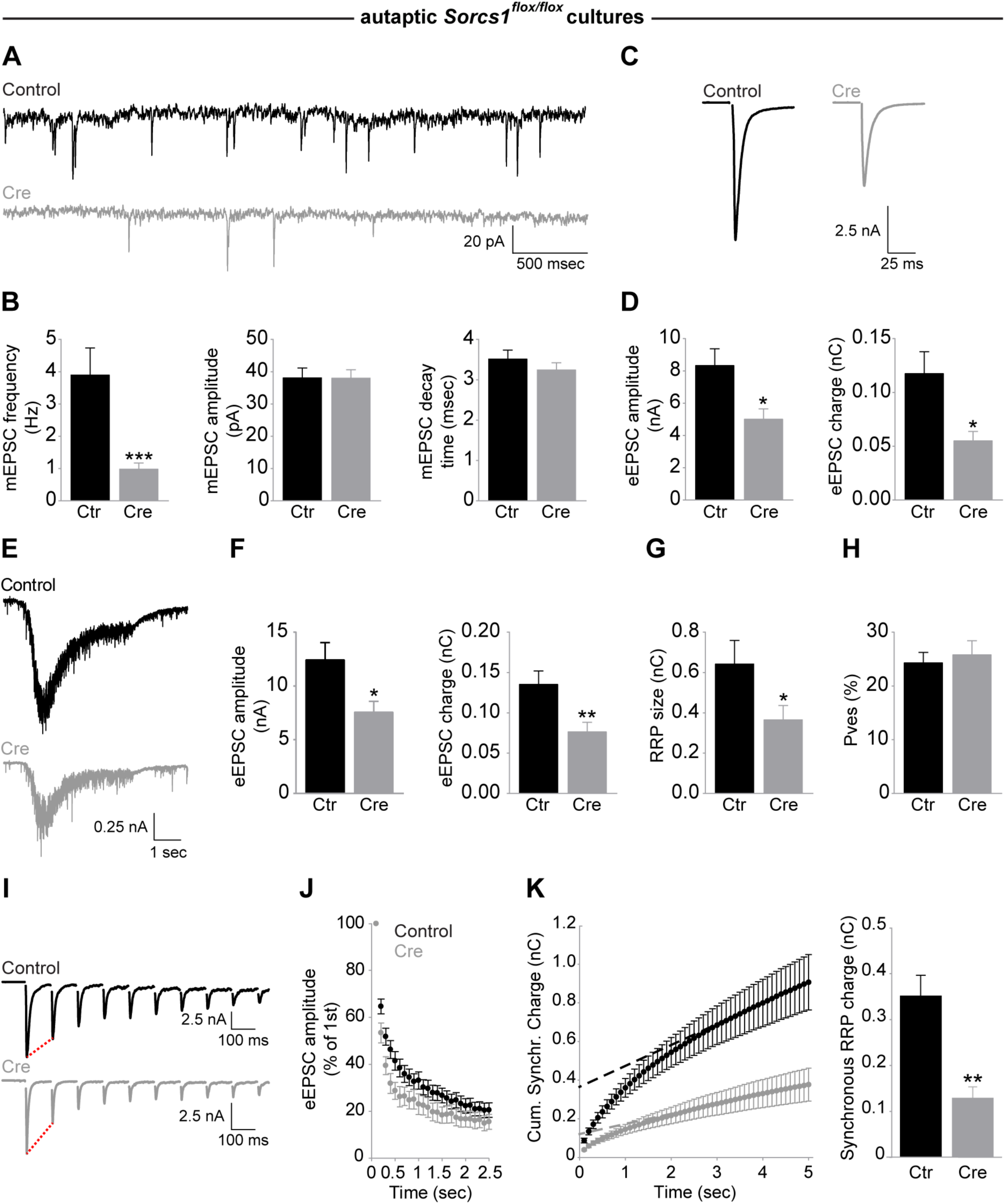
SorCS1 is required for presynaptic function. A, Example traces of mEPSCs recorded from DIV14–DIV16 *Sorcs1^flox/flox^*autaptic cortical neurons electroporated with EGFP (Control) or Cre-EGFP (Cre). B, mEPSC frequency, but not amplitude and decay time, is decreased in *Sorcs1* KO neurons. Control (n = 18 neurons); Cre (n = 15). ***p < 0.001 (Mann-Whitney test, 3 independent experiments). C, Example traces of eEPSCs recorded from DIV14–DIV16 *Sorcs1^flox/flox^*autaptic cortical neurons electroporated with EGFP or Cre-EGFP. D, eEPSC amplitude and eEPSC charge are decreased in *Sorcs1* KO neurons. Control (n = 17 neurons); Cre (n = 13). *p < 0.05 (Mann-Whitney test, 4 independent experiments). E, Example traces of sucrose responses recorded from DIV14 *Sorcs1^flox/flox^* autaptic cortical neurons electroporated with EGFP or Cre-EGFP. F-H, Decreased eEPSC amplitude, eEPSC charge (F) and readily releasable pool (RRP) size (G) in *Sorcs1* KO neurons, but unaltered vesicular release probability (Pves) (H). Control (n = 17 neurons); Cre (n = 21). *p < 0.05; **p < 0.01 (Mann-Whitney test, 3 independent experiments). I, Example traces of train stimulations (10 Hz) recorded from DIV14–DIV16 *Sorcs1^flox/flox^*autaptic cortical neurons electroporated with EGFP or Cre. J, Increased depression during 10 Hz stimulation in *Sorcs1* KO neurons. Control (n = 18 neurons); Cre (n = 14); 3 independent experiments. K, Estimation of the RRP size used during neuronal activity by back-extrapolating a linear fit of the steady state current towards the ordinate axis intercept, which represents the initial RRP size before train stimulations (10 Hz). RRP size of active synapses was reduced in DIV14–DIV16 *Sorcs1* KO neurons. Control (n = 18 neurons); Cre (n = 13). **p < 0.01 (Mann-Whitney test, 3 independent experiments). Graphs show mean ± SEM.

## Discussion

The membrane trafficking mechanisms by which neurons control the polarized distribution and abundance of synaptic receptors are poorly understood. Here, we show that SorCS1-mediated sorting in dendritic endosomes controls a balance between axonal and dendritic surface levels of Nrxn and maintains Nrxn axonal surface polarization required for presynaptic differentiation and function.

### A SorCS1-Rab11FIP5/Rip11 interaction sorts axonal cargo from EEs to REs

The mechanism by which SorCS1 regulates trafficking of neuronal receptors has remained unclear. We find that SorCS1 controls a balance between axonal and dendritic surface polarization of its cargo Nrxn. In mature neurons, newly synthesized Nrxn1α traffics to the somatodendritic surface, followed by endocytosis and transcytosis to the axon. Accordingly, blocking endocytosis, EE-, and RE-mediated transport all result in accumulation of Nrxn1α on the dendritic surface. These effects are mimicked by *Sorcs1* KO. We find that endosomal SorCS1 controls the transition of endocytosed Nrxn1α from EEs to REs and identify the Rab11-interacting protein Rab11FIP5/Rip11 as a novel SorCS1 cytoplasmic interactor. Interference with Rip11 function alters the axonal-dendritic surface balance of Nrxn1α in the same way that *Sorcs1* KO does. Rab11FIPs function as linkers between Rab11 and motor proteins to promote sorting of cargo from EEs to REs [37, 47] and transcytosis in polarized epithelial cells [38, 39], but their function in neuronal transcytosis has not been explored. Our data suggest that SorCS1/Rip11 form a protein complex that localizes to dendritic endosomes and sorts internalized Nrxn1α from early to recycling endosomes, thereby biasing the trafficking of Nrxn1α-containing REs to the axon while preventing Nrxn1α mis-sorting to the dendritic surface and lysosomes.

SorCS1-mediated sorting of Nrxn1α in dendrites can be circumvented by deleting the 4.1-binding motif in the cytoplasmic tail of Nrxn1α, which results in strong polarization of Nrxn1α to the axonal surface. Surface polarization of Nrxn1α Δ4.1 is unaffected by blocking endocytosis or removing *Sorcs1*. It remains to be determined whether Nrxn1α’s 4.1-binding motif, which lacks canonical YxxØ and [D/E]xxx[L/I] dendritic sorting motifs [48] but contains several tyrosine residues, is required for initial sorting to dendrites or for insertion in the dendritic membrane. Binding of the AMPAR subunit GluA1 to the 4.1N protein is required for GluA1 membrane insertion [49]. We find that the extracellular domain of Nrxn1α is required for somatodendritic sorting (data not shown), suggesting that deletion of the 4.1-binding motif in Nrxn1α impairs dendritic insertion but not initial sorting to this compartment.

Our results show that loss of SorCS1 impairs transcytotic trafficking of Nrxn and L1/NgCAM, but not of Caspr2, which traffics via the selective endocytosis/retention pathway, supporting a role for SorCS1 in regulating transcytotic trafficking of axonal cargo proteins. It is unlikely that all membrane proteins that have been identified using surface proteome analysis to depend on SorCS1-3 for their surface trafficking [20,23,24] undergo transcytosis. The SorCS1 cargo amyloid precursor protein (APP) [50–52] however is transcytosed in human neurons [53]. SorCS1 likely interacts with additional proteins, such as the retromer complex [50], to regulate the surface trafficking of a diverse set of cargoes.

### SorCS1-mediated surface polarization of Nrxn is required for presynaptic differentiation and function

The selective impairment in Nlgn1-induced presynaptic differentiation in *Sorcs1* KO neurons and rescue thereof by Nrxn1α Δ4.1, but not by WT Nrxn1α, indicates that this defect is due to a decrease in the abundance of axonal surface Nrxns. Nrxns play key roles in neurotransmitter release [5–7]. Alpha-Nrxn KO in neocortical cultured slices reduces mEPSC/mIPSC frequency and eIPSC amplitude [7]. Beta-Nrxn KO in cortical neurons decreases mEPSC frequency and eEPSC amplitude [5]. Our results show that loss of SorCS1 in autaptic cortical neurons decreases mEPSC frequency, eEPSC amplitude, and sucrose-evoked responses, in line with the reported decreased fraction of active synapses in *Sorcs1* KO neurons [20]. Furthermore, we found an increase in synaptic depression of normalized eEPCSs during stimulus trains, similar to alpha-Nrxn KO neurons [7]. Together, these results suggest that mis-trafficking of Nrxn is a major contributor to the defects in presynaptic function in *Sorcs1* KO neurons. These defects are likely attributed to a decrease in active synapse number and a presynaptic defect in neurotransmitter release, rather than a decrease in excitatory synapse density, which is unaffected in cultured *Sorcs1* KO cortical neurons [20] and in Nrxn TKD hippocampal neurons [46].

The axonal-dendritic balance of Nrxn1α is developmentally regulated. Early in neuronal development, endogenous Nrxn1α is enriched on the axonal surface and traffics directly to the axon after TGN exit. As neurons mature, Nrxn1α follows an indirect trafficking pathway after TGN exit via SorCS1-mediated sorting in dendritic endosomes and transcytosis to the axonal surface. The biological function of this developmental regulation and circuitous trafficking route is not clear. Transcytosis could provide an efficient way of delivering receptors from a reservoir of readily-synthesized proteins. Similar to other transcytosed axonal cargo, a dendritic pool of Nrxn may supply the demand for receptors along the axonal surface, either constitutively [54, 55] or in response to ligand-receptor interactions and signalling [54, 56].

Dendritically localized Nrxns might have a function in this compartment. Postsynaptically expressed Nrxn1 decreases Nlgn1’s synaptogenic effect in hippocampal neurons, likely by *cis*-inhibition of Nlgn1 [32]. In retinal ganglion cells, a shift in Nrxn localization away from dendrites has been proposed to allow dendritic innervation [30]. In cortical neurons however we observe the opposite: Nrxn surface expression in dendrites increases with maturation. Dendritic Nrxns might modulate the function of postsynaptic Nrxn ligands, such as Nlgns, which control postsynaptic neurotransmitter receptor function [57], or LRRTM1, which shapes presynaptic properties [58]. At the *C. elegans* neuromuscular junction, the ectodomain of postsynaptic Nrxn is proteolytically cleaved and binds to the presynaptic α2δ calcium channel subunit to inhibit presynaptic release [59].

In conclusion, impaired trafficking of critical cargo proteins in the absence of SorCS1 and resulting defects in synaptic function might contribute to the pathophysiology of the neurodevelopmental [60–63] and neurodegenerative disorders [52, 64] with which *SORCS1* has been associated. These observations underscore the notion that intracellular sorting is a key contributor to the proper maintenance of synaptic protein composition and function.

## Materials and Methods

### Animals

All animal procedures were approved by the Institutional Animal Care and Research Advisory Committee of the KU Leuven (ECD P015/2013 and ECD P214/2017) and were performed in accordance with the Animal Welfare Committee guidelines of the KU Leuven, Belgium. The health and welfare of the animals was supervised by a designated veterinarian. The KU Leuven animal facilities comply with all appropriate standards (cages, space per animal, temperature, light, humidity, food, water), and all cages are enriched with materials that allow the animals to exert their natural behavior. Both males and females were used for all experiments. To the best of our knowledge, we are not aware of an influence of sex on the parameters analysed in this study. Wistar rats were obtained from Janvier Labs. CD-1 and C57BL/6J mice were obtained from the KU Leuven Mouse Facility. C57BL/6J *Sorcs1^flox/flox^* mice were previously described [50] and kindly provided by Alan D. Attie (University of Wisconsin-Madison, USA).

C57BL/6J *Nrxn1α^HA^* and C57BL/6J *Sorcs1^HA^* knock-in (KI) mice were generated by CRISPR/Cas9-mediated homology directed repair (HDR) targeting the mouse *Nrxn1* locus at exon 2 and *Sorcs1* locus at exon 1. *Sorcs1^HA^*was commercially purchased from Cyagen, whereas *Nrxn1α^HA^* was generated in-house at the Mutamouse core facility (KU Leuven/VIB). To generate *Nrxn1α^HA^*KI, CRISPR/Cas9 components were microinjected in zygotes as ribonucleoproteins (RNPs) composed of: crRNA (guide RNA sequence: 5’-TCCACTGGCCCTCGGCGCCC-3’) (IDT), tracrRNA (IDT, Cat #1072534), ssODN (oligo DNA donor sequence: 5’-TGCTGTCTCCTCTGCCTGTCGCTGCTGCTGCTGGGCTGCTGGGCAGAGCTGGGCAGCGGGTACCCATACGACGTCCCAGATTACGCTCTGGAGTTTCCGGGCGCCGAGGGCCAGTGGACGCG CTTCCCCAAGTGGAACGCGTGCTGC-3’) (IDT) and Cas9 (IDT, Cat #1074182). To generate *Sorcs1^HA^* KI, the donor DNA (5’-GTCGGAGCCGCCGGGACATGCTAAAGGATGGAGGGCAGCAGGGGCTTGGGACTGGCGCATACCCATACGATGTTCCAGATTACGCTCGGGACCCGGACAAAGCCACCCGCTTCCGGATGGAGGAGCTGAGACTGACCAGCACCACA-3’) and guide RNA (5’-TGGGACTGGCGCACGGGACCCGG-3’) were used. Insertion of the HA epitope tag was validated by sequencing, restriction digest (for *Nrxn1α^HA^*) and western blot. To mitigate possible CRISPR/Cas9-mediated off-targets, *Nrxn1α^HA^* and *Sorcs1^HA^*mice were backcrossed to WT C57BL/6J. *Nrxn1α^HA^* and C57BL/6J *Sorcs1^HA^* mice are fertile and viable.

### Cell Lines

HEK293T-17 human embryonic kidney cells (available source material information: fetus) were obtained from American Type Culture Collection (ATCC, Cat #CRL-11268). HEK293T-17 cells were grown in DMEM (Thermo Fisher Scientific, Cat #11965092) supplemented with 10% (vol/vol) FBS (Thermo Fisher Scientific, Cat #10270106) and 1% (vol/vol) penicillin/streptomycin (Thermo Fisher Scientific, Cat #15140122).

### Neuronal Cultures

#### Primary neuronal cultures for imaging

Neurons were cultured from E18-E19 Wistar rat embryos (Janvier labs) (cortical), from E18–E19 WT C57BL/6J mice embryos (cortical and hippocampal), from E18–E19 *Sorcs1^flox/flox^* mice (cortical), from E18–E19 *Sorcs1^HA^* mice (cortical) and E18–E19 *Nrxn1α^HA^* mice (cortical), as previously described [20]. Briefly, dissected hippocampi and cortices were incubated with trypsin [0.25% (vol/vol), 15 min, 37 °C] (Thermo Fisher Scientific, Cat #15090046) in HBSS (Thermo Fisher Scientific, Cat #14175095) supplemented with 10 mM HEPES (Thermo Fisher Scientific, Cat #15630080). After trypsin incubation, hippocampi and cortices were washed with MEM (Thermo Fisher Scientific, Cat #11095080) supplemented with 10% (v/v) horse serum (Thermo Fisher Scientific, Cat #26050088) and 0.6% (wt/vol) glucose (MEM-horse serum medium) three times. The cells were mechanically dissociated by repeatedly pipetting the tissue up and down in a flame-polished Pasteur pipette, and then plated on poly-D-lysine (Millipore, Cat #A-003-E) and laminin (Sigma-Aldrich, Cat #L2020) coated glass coverslips (Glaswarenfabrik Karl Hecht, Cat #41001118) in 60 mm culture dishes (final density: 4 x 10^5^–7 x 10^5^ cells per dish), containing MEM-horse serum medium. Once neurons attached to the substrate, after 2–4 hr, the coverslips were flipped over an astroglial feeder layer in 60-mm culture dishes containing neuronal culture medium: Neurobasal medium (Thermo Fisher Scientific, Cat #21103049) supplemented with B27 (1:50 dilution; Thermo Fisher Scientific, Cat #17504044), 12 mM glucose, glutamax (1:400 dilution; Thermo Fisher Scientific, Cat #35050061), penicillin/streptomycin (1:500 dilution; Thermo Fisher Scientific, Cat #15140122), 25 μM β-mercaptoethanol, and 20 μg/mL insulin (Sigma-Aldrich, Cat #I9278). Neurons grew face down over the feeder layer but were kept separate from the glia by wax dots on the neuronal side of the coverslips. To prevent overgrowth of glia, neuron cultures were treated with 10 μM 5-Fluoro-2′-deoxyuridine (Sigma-Aldrich, Cat #F0503) after 3 d. Cultures were maintained in a humidified incubator of 5% (vol/vol) CO_2_/95% (vol/vol) air at 37 °C, feeding the cells once per week by replacing one-third of the medium in the dish. Cortical neurons from WT and *Sorcs1^flox/flox^*mice were electroporated with DNA just before plating using an AMAXA Nucleofector kit (Lonza, Cat #VPG-1001). Cortical neurons from *Sorcs1^flox/flox^* mice were transfected at DIV7 using Effectene (Qiagen, Cat #301425). Rat cortical neurons, and neurons from WT (cortical and hippocampal), *Sorcs1^flox/flox^* (cortical) and *Nrxn1α^HA^* (cortical) mice were transfected at DIV6–DIV8 using calcium phosphate. Mouse cortical neurons from *Nrxn1α^HA^* and *Sorcs1^HA^*were mixed with WT cortical neurons which allowed for reliable immunodetection of endogenous HA-Nrxn1α and HA-SorCS1.

Co-culture assays were performed as previously described [65]. Briefly, HEK293T cells grown in DMEM (Thermo Fisher Scientific, Cat #11965092) supplemented with 10% (vol/vol) FBS (Thermo Fisher Scientific, Cat #10270106) and 1% (vol/vol) penicillin/streptomycin (Thermo Fisher Scientific, Cat #15140122), were transfected with FLAG-tagged Nlgn1 or GFP-tagged NGL-3 using Fugene6 (Promega, Cat #E2691). Transfected HEK293T cells were mechanically dissociated 24 hr after transfection and co-cultured with DIV10 cortical neurons prepared from *Sorcs1^flox/flox^* mice for 16 hr.

#### Primary autaptic cortical cultures for electrophysiology

For electrophysiology, we used isolated cortical neurons on astrocyte microislands. In short, cortices were dissected from E18 *Sorcs1^flox/flox^* or WT mice and collected in HBSS (Thermo Fisher Scientific, Cat #14175095) supplemented with 10 mM HEPES (Thermo Fisher Scientific, Cat #15630080). Cortical pieces were incubated for 15 min in HBSS supplemented with tripsin 0.25% (vol/vol) (Thermo Fisher Scientific, Cat #15090046) and 10 mM HEPES for 15 min at 37 °C. After trypsin blocking and washing with MEM (Thermo Fisher Scientific, Cat #11095080) supplemented with 10% (v/v) horse serum (Thermo Fisher Scientific, Cat #26050088) and 0.6% (wt/vol) glucose; neurons were dissociated, counted and plated in Neurobasal medium (Thermo Fisher Scientific, Cat #21103049) supplemented with B27 (1:50 dilution; Thermo Fisher Scientific, Cat #17504044), 12 mM glucose, glutamax (1:400 dilution; Thermo Fisher Scientific, Cat #35050061), penicillin/streptomycin (1:500 dilution; Thermo Fisher Scientific, Cat #15140122) and 25 μM β-mercaptoethanol. Dissociated neurons were electroporated (*Sorcs1^flox/flox^*: EGFP or Cre-EGFP; WT: L315 control construct or L315 Nrxn triple knockdown construct) just before plating using AMAXA Nucleofector kit (Lonza, Cat #VPG-1001) and plated at 2500/cm^2^ on micro islands of mouse (CD-1) glia. Glial islands were obtained by first coating glass coverslips with 0.15% (wt/vol) agarose. After drying and UV sterilization (30 min), customized stamps were used to create dots (islands, diameter 200–250 µm) using a substrate mixture containing 0.25 mg/mL rat tail collagen (Corning, Cat #354236) and 0.4 mg/mL poly-D-lysine (Sigma-Aldrich, Cat #P7405) dissolved in 17 mM acetic acid. After drying/UV treatment of the islands, ∼25000 astrocytes were plated per 30 mm glass coverslip in 6-well plates and allowed to form micro-dot islands in DMEM (Thermo Fisher Scientific, Cat #11965092) supplemented with 10% (vol/vol) FBS (Thermo Fisher Scientific, Cat #10270106) for 3–5 d.

### Plasmids

Constructs were all generated using the Gibson Assembly Cloning Kit (New England Biolabs, Cat #E5510S) by inserting the different DNA fragments (PCR-generated, gBlocks or ultramers) in the final vectors digested with the respective restriction enzymes. pCAG-HA-Nrxn1α(-SS4) was kindly provided by Peter Scheiffele (University of Basel, Switzerland). For pFUGW-mCherry, pFUGW-Cre_T2A_mCherry, pFUGW-Cre_T2A_mCherry_T2A_Nrxn1α(-SS4) WT, pFUGW-Cre_T2A_mCherry_T2A_Nrxn1α(-SS4) Δ4.1, pFUGW-Cre-EGFP, pFUGW-Cre-EGFP_T2A_SorCS1^WT^, pFUGW-Cre-EGFP_T2A_SorCS1^Y1132A^, PCR fragments were inserted into pFUGW vector digested with AgeI and EcoRI. For pcDNA3.1-HA-SorCS1cβ a PCR fragment was inserted into pcDNA3.1(+) (Thermo Fisher Scientific, Cat #V79020) digested with EcoRI. For FCK(0.4)GW-Cre-EGFP a PCR fragment was inserted into FCK(0.4)GW vector digested with AgeI and EcoRI. For pIRESneo3-Str-KDEL_SBP-EGFP-Nrxn1α, a PCR fragment was inserted into pIRESneo3-Str-KDEL_SBP-EGFP-GPI (kindly provided by Franck Perez, Institute Curie, France; Addgene plasmid #65294; RRID: Addgene_65294) digested with FseI and PacI. pcDNA4-SorCS1cβ-myc Y1132A, pcDNA3.1-HA-SorCS1cβ Y1132A and GFP-tagged Rab7 T22N were generated by *in vitro* mutagenesis using the QuikChange II Site-Directed Mutagenesis Kit (Agilent, Cat #200522) from pcDNA4-SorCS1cβ-myc (kindly provided by Alan D. Attie, University of Wisconsin-Madison, USA), pcDNA3.1-HA-SorCS1cβ and GFP-tagged Rab7 WT (kindly provided by Casper Hoogenraad, Utrecht University, The Netherlands), respectively. The following constructs were kindly provided by: pIRESneo3-Str-KDEL_TfR-SBP-EGFP and GFP-tagged TfR (Juan S. Bonifacino, National Institute of Health, USA); pEGFP-N3-Rip11 and pEGFP-C1-Rip11(490–653) (Rytis Prekeris, University of Colorado, USA); GFP-tagged Rab11 WT, GFP-tagged Rab11 S25N and GFP-tagged Rab4 WT (Casper Hoogenraad, Utrecht University, The Netherlands); GFP-tagged Rab5 WT and GFP-tagged Rab5 S34N (Ragna Sannerud, KU Leuven, Belgium); GFP-tagged Dynamin1 WT, GFP-tagged Dynamin1 WT K44A and pcDNA3-HA-Caspr2 (Catherine Faivre-Sarrailh, Aix-Marseille University, France); L315 control and L315 Nrxns TKD (Thomas C. Südhof, Stanford University, USA); FLAG-tagged Nlgn1 (Davide Comoletti, Rutgers University, USA); pEGFP-N1-NGL-3 (Eunjoon Kim, Korea Advanced Institute of Science and Technology, South Korea); and myc-tagged L1 (Dan P. Felsenfeld, CHDI Foundation, USA). All DNA constructs used in this study were verified by sequencing. **See Table S1 for a list of plasmids used in this study.**

### Neuron Transfection

Rat cortical neurons (live-cell imaging experiments), and neurons from WT (cortical and hippocampal), *Sorcs1^flox/flox^*(cortical) and *Nrxn1α^HA^* (cortical) mice were transfected at DIV6–DIV8 using calcium phosphate, with the exception of the DIV3 live-cell imaging experiment, in which rat cortical neurons were transfected at DIV2. 2 μg of DNA or 1 μg of DNA from each DNA construct (double co-transfections) were used per coverslip; with the exception of co-transfections of GFP-tagged Rab proteins or Rip11 (WT and dominant negatives) with extracellular HA-tagged Nrxn1α, in which 0.75 μg and 1.25 μg of DNA was used for GFP-tagged and HA-Nrxn1α DNA constructs, respectively. Briefly, DNA plasmids were diluted in Tris-EDTA buffer (10 mM Tris-HCl and 2.5 mM EDTA, pH 7.3), followed by dropwise addition of CaCl_2_ solution (2.5 M CaCl_2_ in 10 mM HEPES, pH 7.2) to the plasmid DNA-containing solution to give a final concentration of 250 mM CaCl_2_. This solution was subsequently added to an equal volume of HEPES-buffered solution (274 mM NaCl, 10 mM KCl, 1.4 mM Na_2_HPO_4_, 42 mM HEPES, pH 7.2) and vortexed gently for 3 s. This mixture, containing precipitated DNA, was then added dropwise to the coverslips in a 12-well plate, containing 250 μL of conditioned neuronal culture medium with kynurenic acid (2 mM), followed by 2 hr incubation in a 37 °C, 5% (vol/vol) CO_2_/95% (vol/vol) air incubator. After 2 hr, the transfection solution was removed, after which 1 mL of conditioned neuronal culture medium with kynurenic acid (2 mM) slightly acidified with HCl (∼5 mM final concentration) was added to each coverslip, and the plate was returned to a 37 °C, 5% (vol/vol) CO_2_/95% (vol/vol) air incubator for 20 min. Finally, coverslips were then transferred to the original dish containing the conditioned culture medium in the 37 °C, 5% (vol/vol) CO_2_/95% (vol/vol) air incubator to allow expression of the transfected constructs. Protein expression was typically for 24 hr (live-cell imaging) or 48 hr (immunocytochemistry).

### Immunocytochemistry

Cells were fixed for 10–15 min in 4% (wt/vol) sucrose and 4% (wt/vol) paraformaldehyde in PBS (PBS: 137 mM NaCl, 2.7 mM KCl, 1.8 mM KH_2_PO_4_ and 10 mM Na_2_HPO_4_, pH 7.4) at room temperature, and permeabilized with PBS + 0.25% (vol/vol) Triton X-100 for 5 min, at 4 °C. Neurons were then incubated in 10% (wt/vol) bovine serum albumin (BSA) in PBS for 1 hr at room temperature to block non-specific staining, and incubated in appropriate primary antibodies diluted in 3% (wt/vol) BSA in PBS (overnight, 4 °C). After washing 3 times in PBS, cells were incubated with the secondary antibodies diluted in 3% (wt/vol) BSA in PBS (1 hr, room temperature). The coverslips were mounted using Prolong Gold Antifade mounting medium (Thermo Fisher Scientific, Cat #P36930).

In all immunocytochemistry experiments performed to determine the subcellular distribution of proteins of interest (Nrxns, SorCS1, Caspr2, L1) the axonal compartment was labeled using antibodies against Ankyrin-G, a marker of the axon initial segment (NeuroMab, Cat #73-146 or Santa Cruz Biotechnology, Cat #sc-31778), and the somatodendritic compartment was labeled using an anti-MAP2 antibody (Abcam, Cat #ab5392). **See Table S2 for a list of antibodies used in this study.**

#### Surface immunostaining of HA-tagged proteins

For surface immunostaining of exogenous HA-Nrxn1α and HA-Caspr2, live mouse cortical and hippocampal neurons were incubated with rabbit anti-HA (1:1000 dilution; Sigma-Aldrich, Cat #H6908) diluted in conditioned neuronal culture medium for 15 min at room temperature. For surface immunostaining of endogenous HA-Nrxn1α, live mouse cortical *Nrxn1α^HA^* KI neurons were incubated with rabbit anti-HA (1:100 dilution; Cell Signaling Technology, Cat #3724) diluted in conditioned neuronal culture medium for 20 min at room temperature. Neurons were then fixed in 4% (wt/vol) sucrose and 4% (wt/vol) paraformaldehyde in PBS for 10 min at room temperature, followed by several washes in PBS and blocking in 10% (wt/vol) BSA in PBS for 1 hr at room temperature. Neurons were then incubated with anti-rabbit secondary antibody diluted in 3% (wt/vol) BSA in PBS (1 hr, room temperature). Following permeabilization, neurons were processed for immunocytochemistry as described above.

#### Surface immunostaining of extracellular myc-tagged L1

To allow surface immunostaining of myc-L1 (live labeling proved to be impossible), neurons were fixed first, followed by several washes and blocking, and then incubated with mouse anti-myc (1:1000 dilution; Santa Cruz Biotechnology, Cat #sc-40) diluted in 3% (wt/vol) BSA in PBS overnight at 4 °C. Subsequently, neurons were incubated with the respective secondary antibody for 1 hr at room temperature, and processed for immunocytochemistry as described above.

#### Antibody pulse-chase experiments

Cultured living neurons were incubated at room temperature for 10 min in the presence of a high concentration (1:250) of mouse anti-HA antibody (Covance, Cat #MMS-101P), against extracellular HA-tagged Nrxn1α and SorCS1, diluted in conditioned medium. Neurons were then washed with pre-warmed PBS at 37 °C to remove the unbound antibody, and were further incubated in antibody free conditioned medium in a 37 °C, 5% (vol/vol) CO_2_/95% (vol/vol) air incubator (for different periods) to allow the internalization of antibody-bound receptors. After this incubation, neurons were fixed in 4% (wt/vol) sucrose and 4% (wt/vol) paraformaldehyde in PBS for 10 min at room temperature. Next, neurons were either exposed to a super-saturating concentration (1:300) of the first of two secondary antibodies, to label the primary antibody-bound surface pool of protein, and/or incubated overnight with anti-mouse Fab fragments [0.25 mg/mL (Santa Cruz Biotechnology, Cat #715-007-003) in 5% (wt/vol) BSA in PBS] to block all primary antibody-bound receptors that were not internalized and/or not labeled by the first secondary antibody. After permeabilization, cells were processed for immunocytochemistry as described above and the pool of internalized receptors was labeled by incubation with the second secondary antibody (1:1000) for 1 hr at room temperature. This strategy allows differential labeling of cell surface and internalized pools of protein.

#### Dynasore treatment

Mouse cortical or hippocampal neurons expressing extracellular HA-tagged Nrxn1α for 30 hr were either treated with 20 µM Dynasore (Sigma-Aldrich, Cat #D7693) or DMSO (vehicle), added to the coverslips in a 12-well plate containing conditioned neuronal culture medium, for 18 hr. Afterwards, surface HA-Nrxn1α was labeled in live neurons as described above.

### Image Analysis and Quantification

All images obtained from immunocytochemistry experiments in fixed cells were captured on a Leica SP8 laser-scanning confocal microscope (Leica Micro-systems). The same confocal acquisition settings were applied to all images taken from a single experiment. Parameters were adjusted so that the pixel intensities were below saturation. Fiji analysis software was used for quantitative imaging analysis. Z-stacked images were converted to maximal intensity projections and thresholded using constant settings per experiment.

#### Quantification of axonal and dendritic immunofluorescence intensity and ratio of axonal/dendritic immunofluorescence intensity

Fluorescence intensity was measured as the sum of integrated intensity in representative portions of axons and dendrites using Ankyrin-G and MAP2 as guides, respectively. Axonal and dendritic intensities were divided by neuritic length and total intensity (axonal + dendritic) (‘*Relative Axonal Intensity’* and ‘*Relative Dendritic Intensity’*) to adjust measurements across cells with varying expression levels (exogenous expression). The A/D ratio was then calculated by dividing the values of axonal and dendritic intensities obtained for every cell (‘*Axonal/Dendritic Intensity’*). A uniformly distributed protein yields an A/D ratio of around 1. A preferentially dendritically localized protein yields an A/D ratio < 1, whereas a preferentially axonally localized protein yields an A/D ratio > 1. To quantify the fluorescence intensity of endogenous Nrxns, axonal and dendritic intensities were not normalized to the total intensity (‘*Axonal Intensity’* and ‘*Dendritic Intensity’*).

#### Polarity index

In some cases polarization of cargo was determined by using the classical polarity index originally described by [66, 67]. This index does not provide a direct measurement of the axonal and dendritic fluorescent intensities, but it allows a quick estimation of the compartmentalized polarization of proteins of interest. One-pixel-wide lines were traced along three dendrites and representative portions of the axon, using MAP2 and Ankyrin-G as guides, in non-thresholded images. The mean intensities (dendritic was averaged from three dendrites) were used to calculate the dendrite:axon (D:A) ‘*Polarity Index’*. D:A = 1, uniform staining; D:A < 1, preferential axonal staining; D:A > 1, preferential dendritic staining.

#### Antibody pulse-chase experiments

Fluorescence intensity was measured as the sum of integrated intensity in representative portions of axons and dendrites using Ankyrin-G and MAP2 as guides, respectively. Intensities of internalized SorCS1 and Nrxn were divided by neuritic length and by the total intensity (surface + internal) (‘*Relative Intensity of Internalized Nrxn’ or ‘SorCS’*) or by the total internalized intensity (axonal + dendritic) (‘*Relative Intensity of Internalized Nrxn’).* Intensities of surface SorCS1 were also divided by neuritic length and by the total intensity (surface + internal) (‘*Relative Intensity of Surface SorCS1’*).

#### Manders coefficient

Manders coefficient was measured by using the Fiji plugin JACoP [68]. Manders coefficient measures the proportion of the signal from channel ‘a’ that coincides with the signal in channel ‘b’ over the total intensity of ‘a’ – M1 coefficient [69].

#### Colocalization of internalized Nrxn with endosomal markers in Sorcs1 KO cells

The colocalization of internalized Nrxn with endosomal markers was evaluated by measuring the density of double-positive puncta for internalized Nrxn and endosomes and by measuring the intensity of Nrxn present in these puncta. Fluorescence intensity was measured as the sum of mean intensity of internalized Nrxn puncta (defined as 0.02 µm^2^-infinite) in representative portions of axons and dendrites (using Ankyrin-G and MAP2 as guides), colocalizing with endosomal markers. Intensities of internalized Nrxn were divided by neuritic length (‘*Nrxn Intensity in Double Positive Puncta’*). Density of internalized Nrxn puncta (defined as 0.02 µm^2^-infinite), colocalizing with endosomal markers, was also divided by neuritic length (‘*Double Positive Puncta Density’*).

### Live-Cell Imaging

For live-cell imaging of fluorescently tagged Nrxn1α- and TfR-positive vesicles, neurons were transfected with a bicistronic expression plasmid encoding Streptavidin-KDEL and SBP-GFP-Nrxn1α or Streptavidin-KDEL and TfR-SBP-GFP, respectively, using the RUSH system [34]. After transfection, neurons were maintained in Neurobasal medium without B27 supplement, because the presence of D-biotin in B27 interferes with the RUSH system, or in Neurobasal medium supplemented with N-2 supplement (Thermo Fisher Scientific, Cat #P36930), which lacks D-biotin. Live-cell imaging was performed at room temperature (∼20°C) in imaging medium (119 mM NaCl, 5 mM KCl, 2 mM CaCl2, 2 mM MgCl2, 30 mM glucose, and 10 mM Hepes, pH 7.4). For long-term live-cell imaging experiments DIV8–DIV10 or DIV3 rat cortical neurons, after 24–31 hr or 25–29 hr of protein expression, respectively, were imaged on a Nikon Eclipse Ti A1R confocal microscope. Images were collected with a 40x oil objective (1.3 NA; Nikon). Focal drift during the experiment was avoided by using the perfect focus system feature of the Nikon system. Laser intensities were kept as low as possible. Time-lapses lasted for 2,5 hr and frames were acquired as z-stacks every 5 min. Synchronous release of Nrxn1α or TfR was induced by application of 40 μM D-biotin (Sigma-Aldrich, Cat #B4501) 10 min after the beginning of the imaging session. Protein synthesis was inhibited by including 20 µg/mL cycloheximide (Sigma-Aldrich, Cat #C4859) in the imaging medium. For short-term live-cell imaging experiments DIV8–DIV10 rat cortical neurons expressing Streptavidin-KDEL and SBP-GFP-Nrxn1α, 21–30 hr after protein expression, were imaged on a Nikon spinning-disk confocal microscope equipped with a 60x oil objective (1.4 NA; Nikon). Time-lapses were performed by sequential capture of 200-ms images of SBP-GFP-Nrxn1α every second for 60–120 s. Focal drift during the experiment was corrected automatically using the autofocus feature of the Nikon system. Neurons were imaged either for 20–34 min or 1–2 hr after exposure to 40 μM D-biotin. In both experiments, a single image of live-stained pan-Neurofascin was taken at the beginning of every imaging session.

#### Live labeling of the axon initial segment (AIS)

Before every live-cell imaging experiment, primary rat cortical cultured neurons were live-labeled with an anti-pan-Neurofascin antibody (NeuroMab, Cat #75-172) to distinguish the axon from the dendrites. Live-cell imaging experiments were performed with rat cortical neurons because live-labeling of the AIS with the anti-pan-Neurofascin antibody did not work in mouse cultured neurons in our hands. Briefly, coverslips with neurons were quickly rinsed in pre-warmed neuronal culture medium. Neurons were incubated with anti-pan-Neurofascin antibody (1/500) diluted in conditioned neuronal culture medium for 10 min in a 37 °C, 5% (vol/vol) CO_2_/95% (vol/vol) air incubator. Coverslips were then quickly washed twice with pre-warmed neuronal culture medium. Finally, neurons were incubated with an Alexa-555 anti-mouse (1/400) (Invitrogen, Cat #A31570) secondary antibody diluted in conditioned neuronal culture medium and incubated for 10 min in a 37 °C, 5% (vol/vol) CO_2_/95% (vol/vol) air incubator. Neurons were quickly rinsed twice with pre-warmed neuronal culture medium and used for live-cell imaging experiments.

#### Analysis of Nrxn1α- and TfR-positive vesicle transport

Dendrites and axon were imaged from the same neuron using spinning disk confocal microscopy. Time-lapses were performed by sequential capture of 200-ms images every second for 60–120 s. Acquisitions were analyzed using an Igor-based software developed by Pieter Vanden Berghe and Valérie Van Steenbergen (Lab. for Enteric NeuroScience and Cell Imaging Core, KU Leuven). Kymographs were created using a segmented line along the axon/dendrite (one dendrite was analysed per cell) from the soma towards the neurite tip, so that anterograde movement occurred from left to right. On the kymographs, single vesicle movement episodes were distinguished as tilted straight lines. Single vesicle pauses were distinguished as straight lines. The number of moving anterograde/retrograde particles in the dendrite and axon was determined manually by drawing lines on top of the trajectories of single particles obtained from the kymographs. Vesicles were defined as motile when showing net displacement of ≥ 2 µm.

### Biochemistry

#### Detection of HA-tagged endogenous SorCS1 and Nrxn1α

Adult brains (*Sorcs1^HA^*, P60 total brain) or (*Nrxn1α^HA^*, three-month-old cortices) from wildtype (+/+), heterozygous (+/Tg) and homozygous (Tg/Tg) KI mice were dissected, snap frozen and stored at −80 °C. The tissue was homogenized in homogenization buffer (20 mM HEPES, pH 7.4; 320 mM sucrose, 5 mM EDTA) supplemented with protease inhibitors (Roche, Cat #11697498001) using a glass Dounce homogenizer. Homogenates were spun at 3000 x g for 15 min at 4 °C. Supernatants were collected and protein quantification was performed with Bio-Rad protein Assay (Cat #500-0006). Samples were analyzed by western blot using a mouse anti-HA antibody (Covance, Cat #MMS-101P; *Sorcs1^HA^*KI) or a rabbit anti-HA antibody (Cell Signaling Technology, Cat #3724; *Nrxn1α^HA^* KI). A total protein stain (Ponceau) was used to assess equal protein loading and transfer to nitrocellulose membrane.

#### Immunoprecipitation of HA-tagged endogenous SorCS1

Cortices from adult *Sorcs1^HA^* KI mouse brains were dissected and homogenized in homogenization buffer (50 mM HEPES pH 7.4, 100 mM NaCl, 2 mM CaCl_2_, 2.5 mM MgCl_2_) supplemented with protease inhibitors (Roche, Cat #11697498001) using a Dounce homogenizer. Cortical homogenates were extracted with 1% CHAPSO (VWR, Cat #A1100.0005) in homogenization buffer while rotating end-over-end for 2 hr at 4°C and centrifuged at 100,000 x g for 1 hr at 4°C to pellet insoluble material. Supernatants were precleared by adding 100 µL of Protein-G agarose beads (Thermo Fisher Scientific, Cat #22852BR) and rotating for 1 hr at 4°C, then incubated with 5 µg of mouse IgG or 50 µL of anti-HA magnetic beads (Thermo Fisher Scientific, Cat #88836) and rotated end-over-end overnight at 4°C. 60 µL of Protein-G agarose beads were added to the IgG control sample and rotated for 1 hr at 4°C, followed by three washes in cold extraction buffer and once in PBS. Anti-HA magnetic beads were washed according to the manufacturer’s protocol using a magnetic stand. Protein-G agarose beads and anti-HA magnetic beads were heated for 15 minutes at 40°C in 50 µL 2X sample buffer and analysed by western blotting.

### Lentivirus Production

Second generation VSV.G pseudotyped lentiviruses were produced as described [70, 71]. HEK293T cells were transfected with control (mCherry) or Cre-T2A-mCherry-containing pFUGW vector plasmids and helper plasmids PAX2 and VSVG using Fugene6 (Promega, Cat #E2691). Supernatant was collected 65 hr after transfection and filtered through a 0.45 µm filter (Thermo Fisher Scientific, Cat #723-2545), aliquoted and stored at −80 °C.

### Electrophysiology

Neurons were recorded on DIV11–DIV16. The patch pipette solution contained (in mM): 136 KCl, 18 HEPES, 4 Na-ATP, 4.6 MgCl_2_, 4 K_2_-ATP, 15 Creatine Phosphate, 1 EGTA and 50 U/ml Phospocreatine Kinase (300 mOsm, pH 7.30). The external medium used contained the following components (in mM): 140 NaCl, 2.4 KCl, 4 CaCl_2_, 4 MgCl_2_, 10 HEPES, 10 Glucose (300 mOsm, pH 7.30). Cells were whole-cell voltage clamped at −70 mV with a double EPC-10 amplifier (HEKA Elektronik) under control of Patchmaster v2×32 software (HEKA Elektronik). Currents were low-pass filtered at 3 kHz and stored at 20 kHz. Patch pipettes were pulled from borosilicate glass using a multi-step puller (P-1000; Sutter Instruments). Pipette resistance ranged from 3 to 5 MΩ. The series resistance was compensated to ∼75%. Only cells with series resistances below 15 MΩ were included for analysis. All recordings were made at room temperature. Spontaneous glutamatergic release was (sEPSC) was recorded at −70 mV. Evoked release was induced using brief depolarization of the cell soma (from 70 to 0 mV for 1 ms) to initiate action potential-dependent glutamatergic release (eEPSCs). A fast local multi-barrel perfusion system (Warner SF-77B, Warner Instruments) was used determine the RRP size using external recording solution containing 500 mM sucrose. A custom analysis procedure in Igor Pro (Wavemetrics Inc.) was used for offline analysis of evoked and sucrose responses. Spontaneous events were detected using Mini Analysis program (Synaptosoft).

### Statistical Analysis

Results are shown as average or as average ± SEM, with n referring to the number of analyzed neurons for each group. For most experiments at least 3 independent cultures were included for analysis. Datasets were tested either using Mann-Whitney *U* test or Kruskal-Wallis test by Dunn’s multiple comparisons test. Statistical testing was performed using GraphPad Prism (GraphPad Software). In all instances, statistical significance was defined as follows: n.s. - not significant (p > 0.05), * p< 0.05, ** p < 0.01, *** p < 0.001.

## Acknowledgements

We thank Anders Nykjaer, Pierre Vanderhaeghen, Sara Calafate, Heather Rice, Elsa Lauwers, Ragna Sannerud and Lucía Chávez-Gutiérrez for critical reading of the manuscript and de Wit lab members for discussion and comments. We thank Casper Hoogenraad for experimental advice and reagents, and Ragna Sannerud, Catherine Faivre-Sarrailh, Rytis Prekeris, Juan S. Bonifacino, Jeremy Tavare, Lorna Hodgson, Dan P. Felsenfeld, Takeshi Sakurai, Marco Arese, Thomas C. Südhof, Peter Scheiffele, Thomas Biederer, Eunjoon Kim, Gopal Thinakaran, Ryohei Iwata, Aaron Bowen and Matthew Kennedy for reagents. We thank Lutgarde Serneels for advice on CRISPR/Cas9-based KI mouse generation, and Pieter Vanden Berghe and Valérie Van Steenbergen for live-cell imaging analysis advice and sharing Igor-based software. L.F.R. is supported by Marie Sklodowska-Curie postdoctoral fellowship H2020-MSCA-IF-2014 and FWO Postdoctoral fellowship 12N0316N/12N0319N. B.V. is supported by FWO PhD fellowship 11A0419N. J.d.W. is supported by ERC Starting Grant (#311083); FWO Odysseus Grant; FWO Project grants G094016N and G0C4518N, FWO EOS grant G0H2818N; a Methusalem grant of KU Leuven/Flemish Government, and ERA-NET NEURON SynPathy 2015.

## Author Contributions

L.F.R. and J.d.W. conceived the project and designed experiments. L.F.R. and B.V. conducted and analyzed experiments. J.N. performed characterization of *Sorcs1^HA^*KI mice. K.M.V. assisted with molecular cloning and construct validation. K.D.W. performed electrophysiology experiments on autaptic neurons. L.F.R., B.V., J.N., K.D.W. and J.d.W. analyzed data. L.F.R. and J.d.W. wrote the paper, with input from all authors.

## SUPPLEMENTARY FIGURES and FIGURE LEGENDS

**Figure S1.**
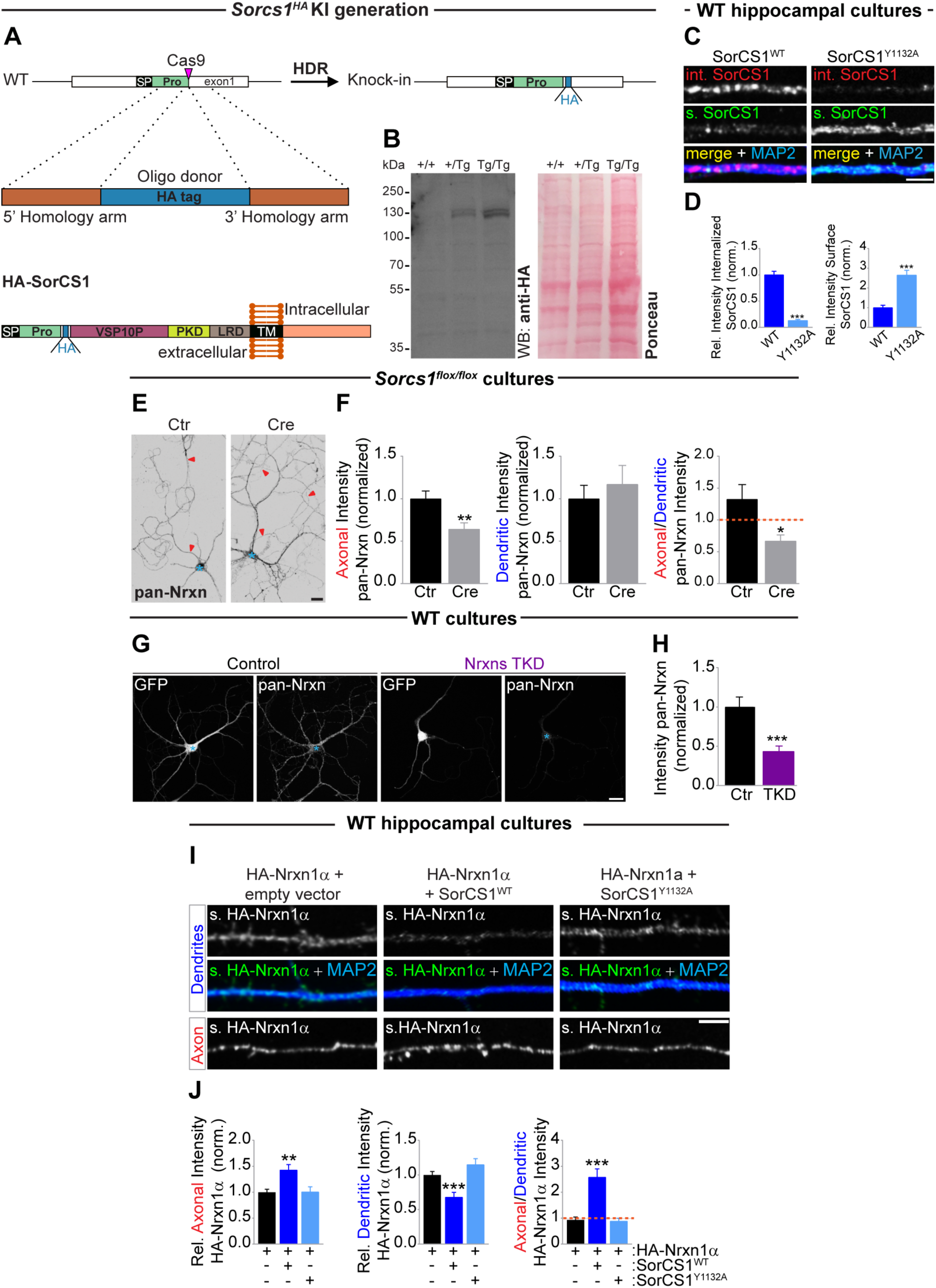
*Sorcs1^HA^* KI mouse generation and SorCS1 regulates axonal surface polarization of Nrxn1α. A, CRISPR/Cas9-mediated generation of *Sorcs1^HA^* KI mice. Orange boxes represent the left and right homology arms. Blue box represents the single-strand DNA (ssDNA) donor oligonucleotide containing the HA-tag. Schematic representation of SorCS1 protein domain organization is shown to illustrate the HA-tagging of HA-SorCS1 downstream of the second furin cleavage site, right before the VPS10P domain (at the amino acid position 144). Domain abbreviations: SP, signal peptide; Pro, pro-peptide; VPS10P, vacuolar protein sorting 10 protein; PKD, polycystic kidney disease domain; LRD, leucine-rich domain; TM, transmembrane. B, Detection of HA-SorCS1 by western blot in total brain extracts prepared from *Sorcs1^HA^* KI mice (P60). Total protein staining (Ponceau) shows equal loading between lanes. C, High-zoom images of dendritic internalized (int.) and surface (s.) SorCS1 from *DIV*9 WT mouse hippocampal neurons expressing HA-tagged WT SorCS1cβ (WT) or Y1132A for 48 hr. Live neurons were incubated with an anti-HA antibody and pulse-chased for 20 min. Neurons were immunostained for surface HA-SorCS1 (grayscale and green), internalized SorCS1 (grayscale and red) and MAP2 (blue). D, Quantification of (C): internalized SorCS1 fluorescence intensity relative to total levels and normalized to cells expressing SorCS1^WT^, surface SorCS1 fluorescence intensity relative to total levels and normalized to cells expressing WT-SorCS1. WT (n = 28 neurons); Y1132A (n = 30). ***p < 0.001 (Mann-Whitney test, 3 independent experiments). E, High-zoom images of Nrxn1α surface distribution from *DIV*8–*DIV*9 WT mouse hippocampal neurons co-expressing extracellular HA-tagged Nrxn1α and an empty vector, HA-Nrxn1α and WT SorCS1-myc or HA-Nrxn1α and an endocytosis-defective mutant of SorCS1cβ-myc (Y1132A) for 48 hr. Neurons were immunostained for surface HA-Nrxn1α (grayscale and green), MAP2 (blue); Ankyrin-G and SorCS1-myc (not shown). F, Quantification of (E): surface HA-Nrxn1α fluorescence intensity in axon and dendrites relative to total surface levels and normalized to cells expressing the empty vector and ratio of axonal/dendritic surface HA intensity. **p < 0.01; ***p < 0.001 (Kruskal-Wallis test followed by Dunn’s multiple comparisons test, 4 independent experiments; n = 40 neurons for each group). G, *DIV*10 *Sorcs1^flox/flox^* cortical neurons electroporated with EGFP (Ctr) or Cre-EGFP immunostained for pan-Nrxn (grayscale), MAP2, Ankyrin-G and GFP (not shown). H, Quantification of (G): pan-Nrxn fluorescence intensity in axon and dendrites normalized to cells expressing EGFP and ratio of axonal/dendritic pan-Nrxn intensity. Ctr (n = 30 neurons); Cre (n = 29 neurons). *p < 0.05; **p < 0.01 (Mann-Whitney test, 3 independent cultures). Graphs show mean ± SEM. Scale bar, 5 µm (C, I), 20 µm (E, G).

**Figure S2.**
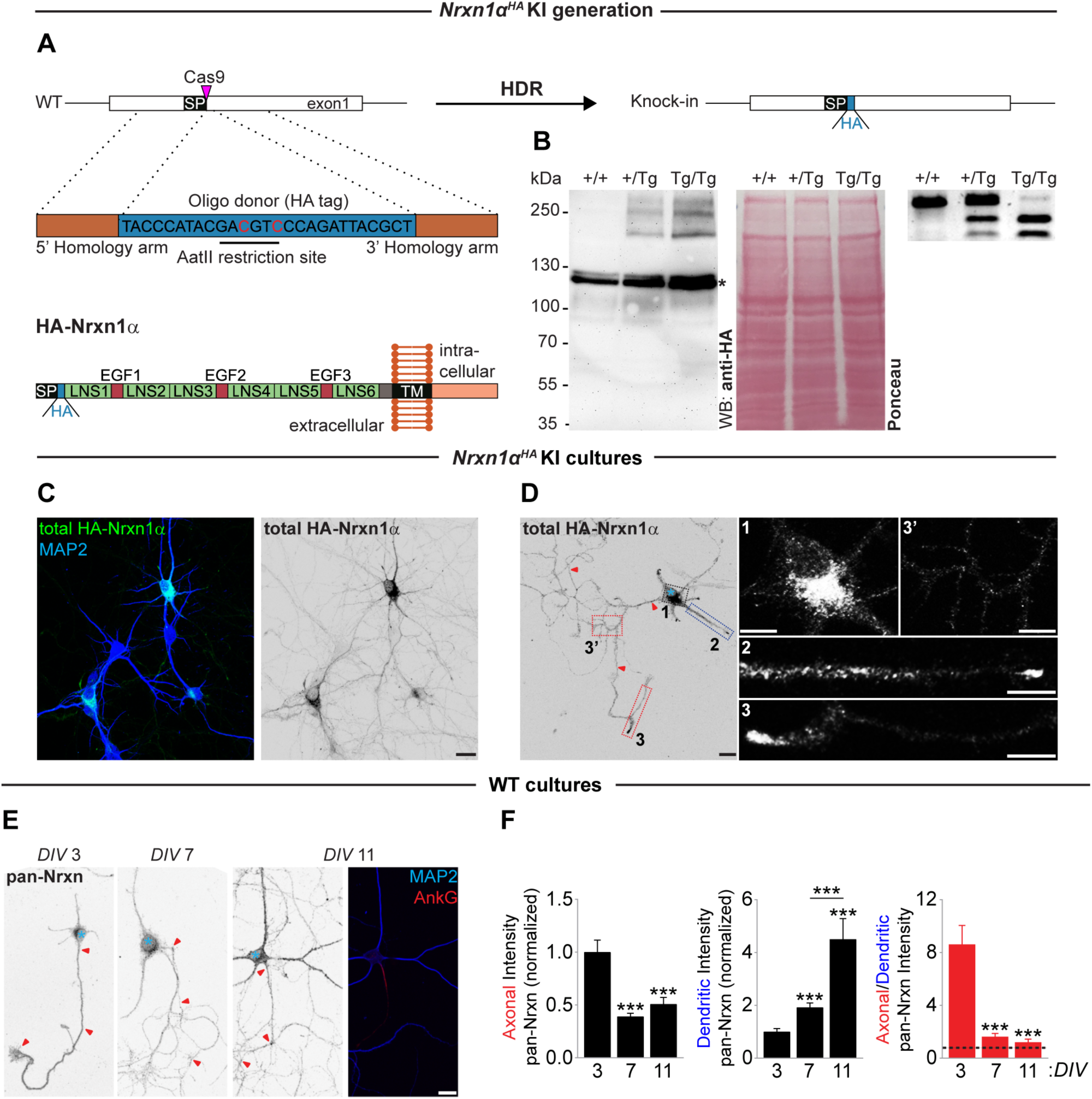
*Nrxn1α^HA^* KI mouse generation and Nrxn1α localizes to the somato-dendritic and axonal compartment. A, Orange boxes represent the left and right homology arms. Blue box represents the single-strand DNA (ssDNA) donor oligonucleotide containing the HA-tag. CRISPR/Cas9-mediated homology directed repair (HDR) allowed for precise HA-tagging of the *Nrxn1* locus right after the signal peptide (SP). An AatII restriction site was introduced in the DNA sequence coding for the HA-tag, by taking advantage of the redundancy of the genetic code, in order to facilitate the distinction between homozygous and heterozygous *Nrxn1α^HA^* mice. Schematic representation of Nrxn1α protein domain organization is shown to illustrate the HA-tagging of HA-Nrxn1α right after the SP in the extracellular domain. Domain abbreviations: LNS, laminin/neurexin/sex hormone-binding globulin; EGF, epidermal growth factor-like; TM, transmembrane. B, Left panels: detection of HA-Nrxn1α by western blot in cortical extracts prepared from adult *Nrxn1α^HA^* KI mice (three months old). Total protein staining by using Ponceau method was used as loading control. Asterisk indicates an aspecific band. Right panel: identification of WT (+/+), heterozygous (+/Tg) and homozygous (Tg/Tg) *Nrxn1α^HA^* mice by generation of a PCR fragment of 499 bp followed by AatII-mediated restriction endonuclease reaction generating two DNA fragments of 312 bp and 187 bp. C and D, Representative images of permeabilized *DIV*7 *Nrxn1α^HA^* mouse cortical neurons cultured together with WT mouse cortical neurons and immunostained for HA-Nrxn1α (green and grayscale), MAP2 (blue) and Ankyrin-G (not shown). Red arrowheads indicate the axon and blue asterisk marks the cell body. High-zoom images of the subcellular distribution of endogenous HA-Nrxn1α: cell body (1), dendrites (2) and axon (3 and 3’). E, Permeabilized *DIV*3, *DIV*7 and *DIV*11 WT mouse cortical neurons labeled with a pan-Nrxn antibody (grayscale). MAP2 (blue) and Ankyrin-G (AnkG; red) identify somatodendritic and axonal compartments, respectively. F, Quantification of (E): pan-Nrxn fluorescence intensity in axon and dendrites normalized to *DIV*3 neurons and ratio of axonal/dendritic pan-Nrxn intensity. *DIV*3 (n = 30 neurons); *DIV*7 (n = 30); *DIV*11 (n = 29). ***p < 0.001 (Kruskal-Wallis test followed by Dunn’s multiple comparisons test, 3 independent cultures). G, Representative images of *DIV*8, *DIV*10 WT mouse cortical neurons electroporated with GFP tagged L315 control construct (Control) or L315 Nrxn triple knockdown construct (Nrxns TKD)^1,2^ immunostained for pan-Nrxn (grasycale) and GFP (grasycale). Blue asterisk marks the cell body. H, Quantification of (G): Nrxn fluorescence intensity normalized to cells expressing GFP (n = 45 neurons for each group). ***p < 0.001 (Mann-Whitney test, 3 independent experiments). Graphs show mean ± SEM. Scale bars, 20 µm (C, D, E); 10 µm (D [high-zoom]).

**Figure S3.**
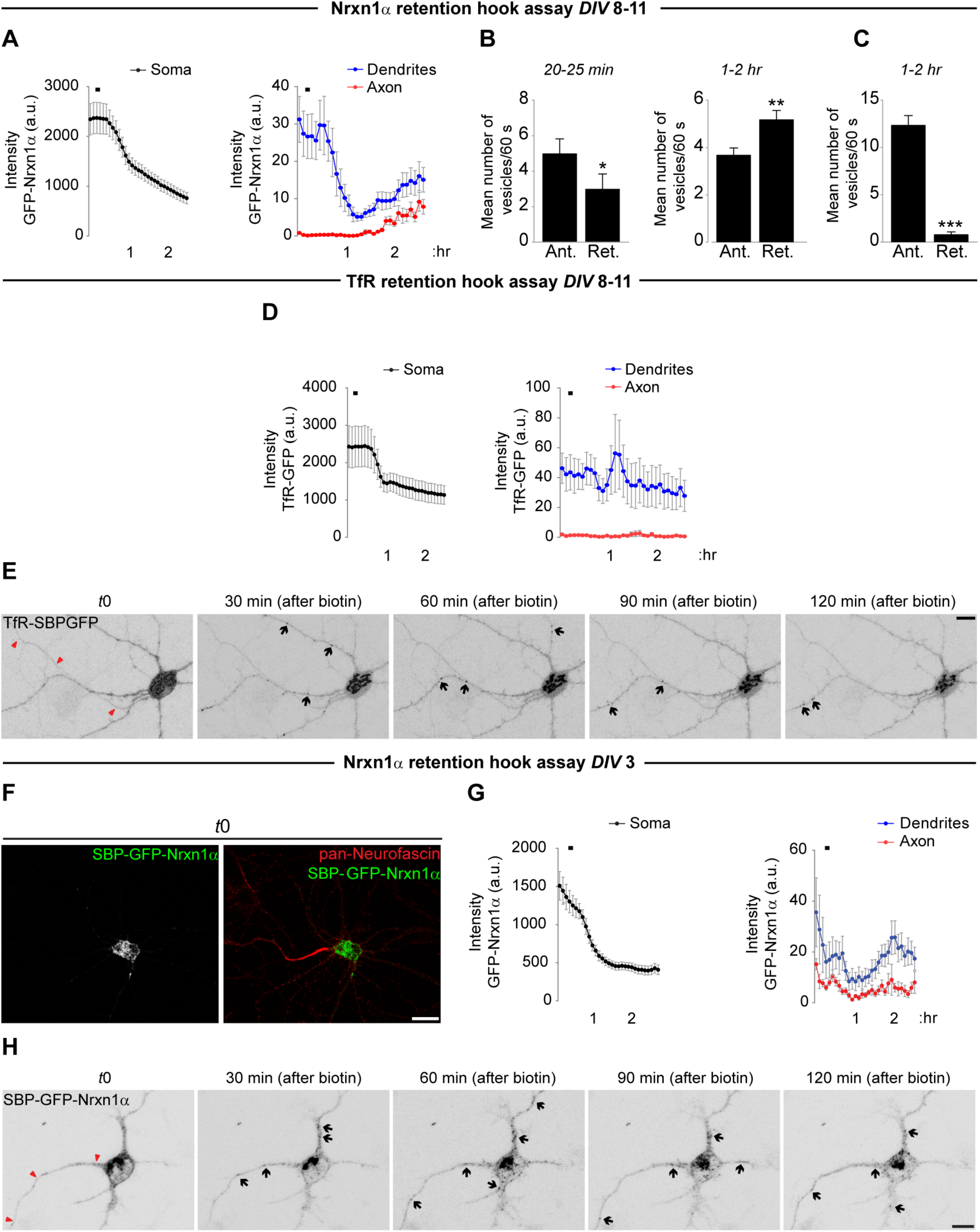
Delayed axonal trafficking of Nrxn1α in mature neurons. A, Quantification of SBP-GFP-Nrxn1α (n = 23 neurons) fluorescence intensity in soma, dendrites and axon in 3 independent experiments. B and C, Mean number of SBP-GFP-Nrxn1α vesicles moving in anterograde and retrograde direction in (B) dendrites and (C) axons. In dendrites, analysis was performed at 2 different time intervals: 20– 25 min, and 1–2 hr after adding biotin; in axons analysis was performed 1–2 hr after biotin. *p < 0.05; **p < 0.01; ***p < 0.001 (Mann-Whitney test). D, Quantification of TfR-SBP-GFP (n = 13 neurons) fluorescence intensity in soma, dendrites and axon in 3 independent experiments. E, Live-cell imaging in *DIV*8–*DIV*10 WT rat cortical neurons co-expressing TfR-SBP-GFP and ER hook. After 24–31 hr of expression neurons were imaged every 5 min for 2.5 hr. Biotin was added 10 min after the beginning of the imaging session. Shown are representative images of TfR-SBP-GFP fluorescence in dendrites and axon before (t0) and 30, 60, 90 and 120 min after adding biotin. Red arrowheads indicate axon and black arrows indicate TfR-SBP-GFP-positive puncta. F, Illustration of the strategy used during live-cell imaging experiments to label the axon. Representative image of a live *DIV*8 WT rat cortical neuron co-expressing SBP-GFP-Nrxn1α (grayscale and green) and streptavidin-KDEL immunostained for pan-Neurofascin (axon initial segment marker, grayscale and red). G, Quantification of GFP-Nrxn1α fluorescence intensity in the soma, dendrites and axons (n = 9 neurons) in 2 independent experiments. H, Live-cell imaging in *DIV*3 WT rat cortical neurons co-expressing SBP-GFP-Nrxn1α and streptavidin-KDEL. After 25–29 hr of expression neurons were imaged every 5 min for 2.5 hr. Biotin was added 10 min after the beginning of the imaging session. See also Supplementary Movie 4. Shown are representative images of GFP-Nrxn1α endogenous fluorescence in dendrites and axons before (t0), 30, 60, 90 and 120 min after adding biotin. Red arrowheads indicate the axon and black arrows indicate GFP-Nrxn1α-positive puncta. Graphs show mean ± SEM. Scale bars, 10 µm (E, F, H).

**Figure S4.**
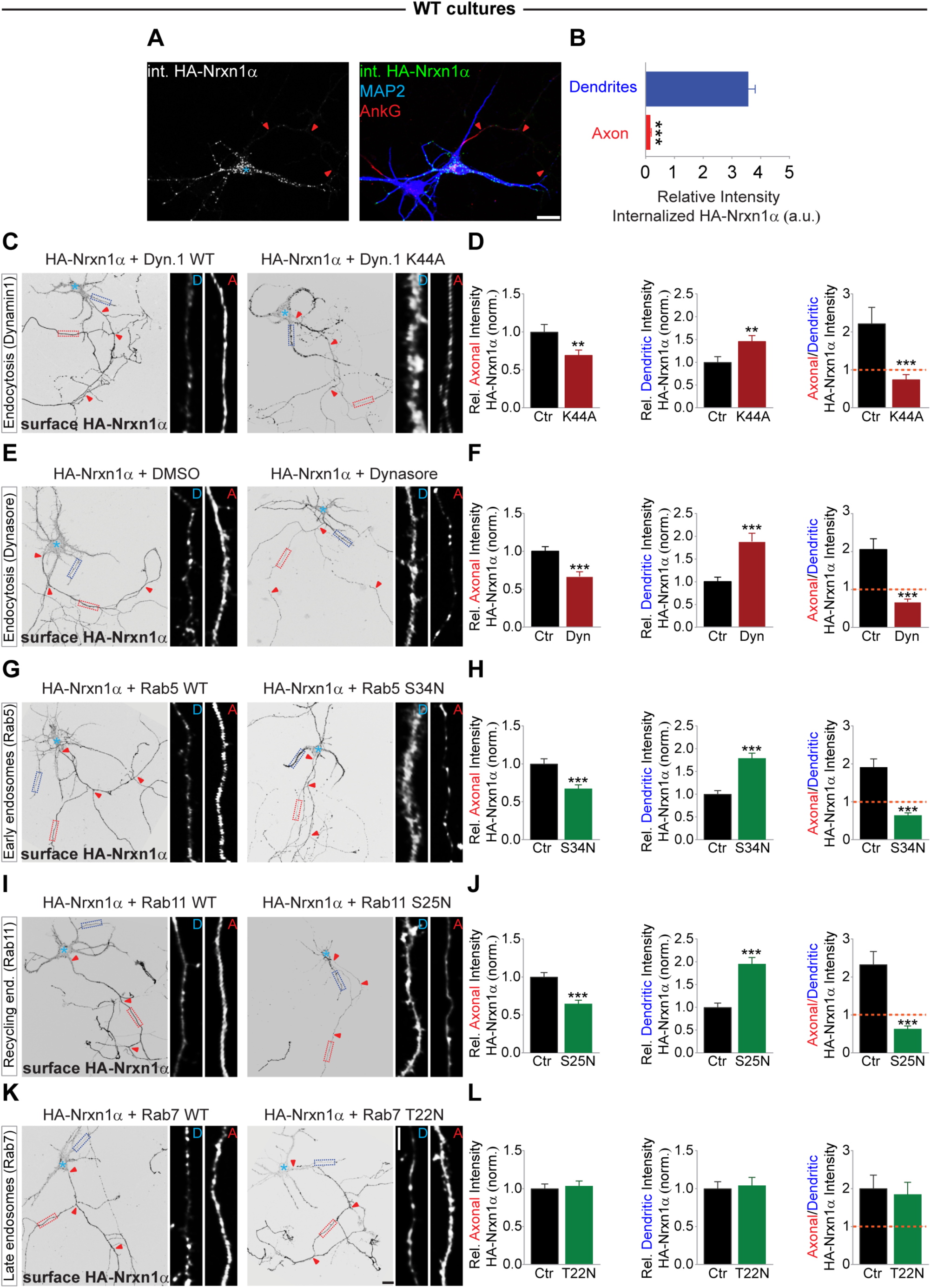
Endocytosis and transport via endosomes are required to sustain axonal levels of Nrxn1α. A, Representative images of mouse cortical neurons transfected with extracellular HA-tagged Nrxn1α. After 48 hr, neurons were live labeled with anti-HA antibody and pulse-chased for 15 min, followed by immunostaining for internalized HA-Nrxn1α (grayscale and green), MAP2 (blue) and Ankyrin-G (red). Red arrowheads indicate the axon and blue asterisk marks the cell body. B, Quantification of internalized Nrxn1α fluorescence intensity in dendrites and axons relative to total levels (n = 15 neurons). ***p < 0.001 (Mann-Whitney test, 3 independent experiments). C, Representative images of *DIV*8–*DIV*10 WT mouse cortical neurons co-expressing HA-Nrxn1α and GFP-tagged WT Dynamin1 (Ctr) or a dominant negative of GFP-Dynamin1 (K44A, to block Dynamin-dependent endocytosis) for 48 hr. Neurons were immunostained for surface HA-Nrxn1α in grayscale; MAP2, Ankyrin-G and GFP (not shown). Red arrowheads indicate the axon and blue asterisk marks the cell body. High-zoom images of dendritic [D, dotted blue box] and axonal [A, dotted red box] Nrxn1α are shown next to the whole cell images. D, Quantification of (C): surface HA-Nrxn1α fluorescence intensity in axon and dendrites relative to total surface levels and normalized to cells expressing Dyn.1 WT and ratio of axonal/dendritic surface HA intensity. Ctr (n = 30 neurons); K44A (n = 30 neurons). E, Representative images of *DIV*8–*DIV*9 WT mouse cortical neurons expressing extracellular HA-tagged Nrxn1α and treated either with DMSO (vehicle, Ctr) or with Dynasore (Dyn) 30 hr after transfection. Neurons were immunostained 18 hr after treatment for surface HA-Nrxn1α (grayscale); MAP2 and Ankyrin-G (not shown). F, Quantification of (E); Ctr (n = 30 neurons); Dyn (n = 27 neurons). G, Representative images of *DIV*8–*DIV*10 WT cortical neurons co-expressing HA-Nrxn1α and GFP-tagged WT Rab5 (Ctr) or a dominant negative of GFP-Rab5 (S34N, to prevent the formation of early endosomes) for 48 hr. Neurons were immunostained for surface HA-Nrxn1α (grayscale); MAP2, Ankyrin-G and GFP (not shown). H, Quantification of (G); Ctr (n = 29 neurons); S34N (n = 27 neurons). I, Representative images of *DIV*8–*DIV*10 WT cortical neurons co-expressing HA-Nrxn1α and GFP-tagged WT Rab11 (Ctr) or a dominant negative of GFP-Rab11 (S25N, to prevent the formation of recycling endosomes) for 48 hr. Neurons were immunostained for surface HA-Nrxn1α (grayscale); MAP2, Ankyrin-G and GFP (not shown). J, Quantification of (I); Ctr (n = 30 neurons); S25N (n = 27 neurons). K, Representative images of *DIV*8–*DIV*10 WT mouse cortical neurons co-expressing extracellular HA-tagged Nrxn1α and GFP-tagged WT Rab7 (Ctr) or a dominant negative of GFP-Rab7 (T22N, to prevent the formation of late endosomes) for 48 hr. Neurons were immunostained for surface HA-Nrxn1α (grayscale); MAP2, Ankyrin-G and GFP (not shown). L, Quantification of (K); Ctr (n = 30 neurons); T22N (n = 29 neurons). *p < 0.05; **p < 0.01; ***p < 0.001 (Mann-Whitney test, 3 independent cultures). Graphs show mean ± SEM. Scale bars, 10 µm (A), 20 µm (C, E, G, I, K); 5 µm (C, E, G, I, K [high-zoom]).

**Figure S5.**
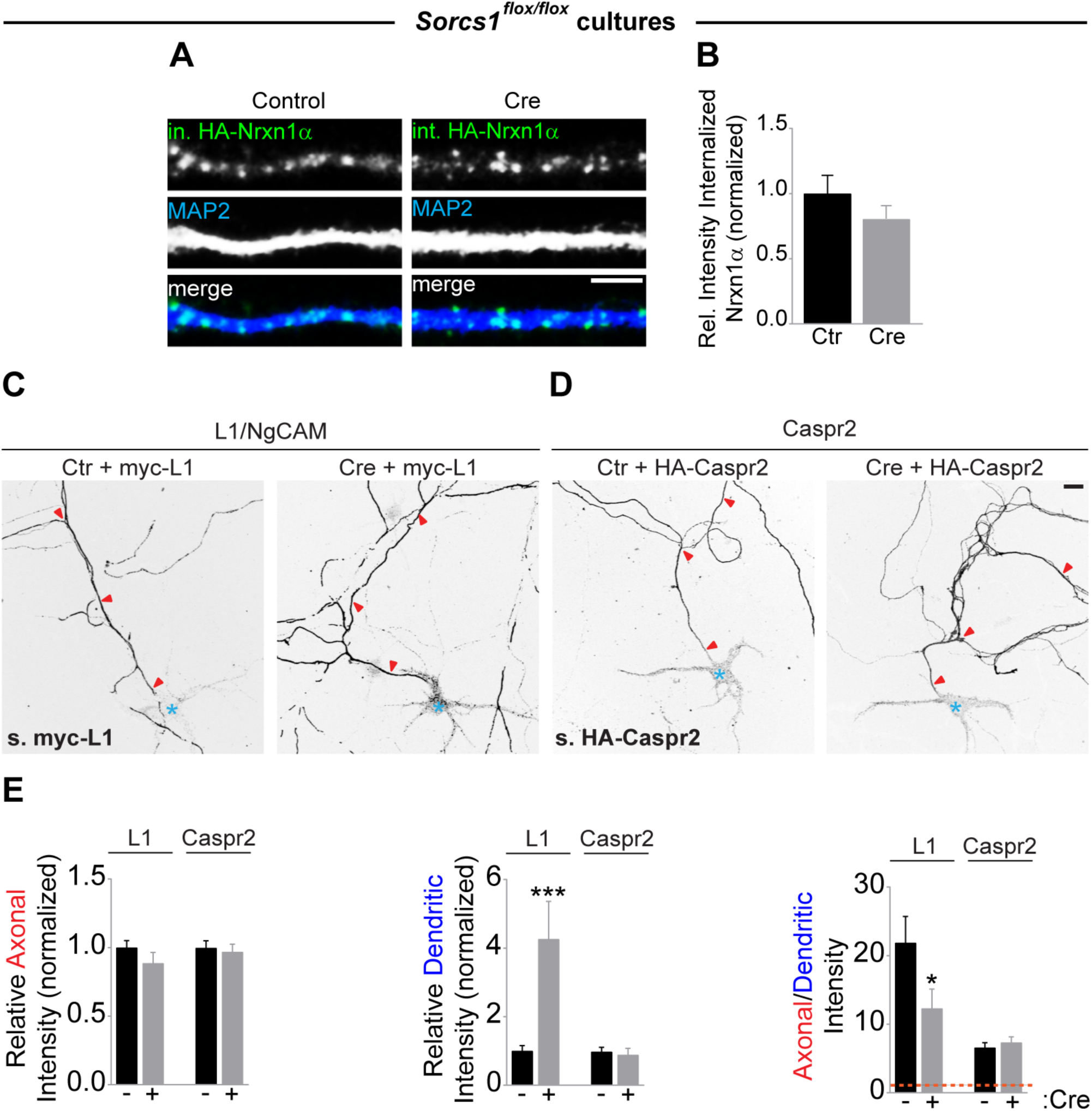
Selective mis-sorting of axonal cargo in the absence of SorCS1. A, High-zoom representative images of dendritic internalized Nrxn1α from *DIV*8, *DIV*10 *Sorcs1^flox/flox^*cortical neurons electroporated with EGFP (Control) or Cre-EGFP (Cre) and transfected with extracellular HA-tagged Nrxn1α. After 48 hr, neurons were live labeled with anti-HA antibody and pulse-chased for 20 min, followed by immunostaining for internalized HA-Nrxn1α (grayscale and green), MAP2 (grayscale and blue), surface HA-Nrxn1α and GFP (not shown). B, Quantification of (A): internalized Nrxn1α fluorescence intensity relative to total levels and normalized to cells expressing GFP (3 independent experiments). Control (n = 30 neurons); Cre (n = 28 neurons). C and D, Representative images of *DIV*8–*DIV*10 *Sorcs1^flox/flox^* mouse cortical neurons electroporated with EGFP (Ctr) or Cre-EGFP, and transfected with myc-L1 (C) or HA-Caspr2 (D) for 48 hr, followed by immunostaining for surface myc-L1 or HA-Caspr2 in grasycale; MAP2, Ankyrin-G and GFP (not shown). E, Quantification of (C and D): surface myc-L1 and Ha-Caspr2 fluorescence intensity in axon and dendrites relative to total surface levels and normalized to cells expressing EGFP and ratio of axonal/dendritic surface L1 and Caspr2 intensity. Ctr_L1 (n = 29 neurons); Cre_L1 (n = 26 neurons); Ctr_Caspr2 (n = 28 neurons); Cre_Caspr2 (n = 25 neurons). *p < 0.05; **p < 0.01; ***p < 0.001 (Mann-Whitney test, at least 3 independent cultures). Graphs show mean ± SEM. Scale bars, 5 µm (A), 20 µm (C, D).

## Supplementary movie captions

**S1 movie**

*DIV*8 WT rat cortical neuron co-expressing the ER-hook (Streptavidin-KDEL) and SBP-GFP-Nrxn1α (grayscale, inverted for clarity) and live-stained for the AIS marker Neurofascin to label the axon. Biotin was added 10 min after the beginning of the imaging session. Cell was recorded every 5 min for 2,5 hr. The axon is indicated. Frame rate: 2 fps.

**S2 movie**

*DIV*9 WT rat cortical neuron co-expressing the ER-hook (Streptavidin-KDEL) and SBP-GFP-Nrxn1α (grayscale, inverted for clarity) and live-stained for the AIS marker Neurofascin to label the axon. Cell was incubated with biotin for 22 min and then recorded every second for 120 s. The axon is indicated. Frame rate: 4 fps.

**S3 movie**

*DIV*10 WT rat cortical neuron co-expressing the ER-hook (Streptavidin-KDEL) and SBP-GFP-Nrxn1α (grayscale, inverted for clarity) and live-stained for the AIS marker Neurofascin to label the axon. Cell was incubated with biotin for 1 hr and 55 min and then recorded every second for 120 s. The axon is indicated. Frame rate: 4 fps.

**S4 movie**

*DIV*9 WT rat cortical neuron co-expressing the ER-hook (Streptavidin-KDEL) and TfR-SBP-GFP (grayscale, inverted for clarity) and live-stained for the AIS marker Neurofascin to label the axon. Biotin was added 10 min after the beginning of the imaging session. Cell was recorded every 5 min for 2,5 hr. The axon is indicated. Frame rate: 2 fps.

**S5 movie**

*DIV*10 WT rat cortical neuron co-expressing the ER-hook (Streptavidin-KDEL) and SBP-GFP-Nrxn1α (grayscale, inverted for clarity) and live-stained for the AIS marker Neurofascin to label the axon. Cell was recorded every second for 30 s. The axon is indicated. Frame rate: 4 fps.

**S6 movie**

*DIV*3 WT rat cortical neuron co-expressing the ER-hook (Streptavidin-KDEL) and SBP-GFP-Nrxn1α (grayscale, inverted for clarity) and live-stained for the AIS marker Neurofascin to label the axon. Biotin was added 10 min after the beginning of the imaging session. Cell was recorded every 5 min for 2,5 hr. The axon is indicated. Frame rate: 2 fps.

